# Revisiting Ancient Polyploidy in Leptosporangiate Ferns

**DOI:** 10.1101/2022.03.12.484015

**Authors:** Hengchi Chen, Yuhan Fang, Arthur Zwaenepoel, Sanwen Huang, Yves Van de Peer, Zhen Li

**Affiliations:** Department of Plant Biotechnology and Bioinformatics, Ghent University, 9052 Ghent, Belgium; VIB Center for Plant Systems Biology, VIB, 9052 Ghent, Belgium; Laboratory for Lingnan Modern Agriculture, Genome Analysis Laboratory of the Ministry of Agriculture and Rural Affairs, Agricultural Genomics Institute at Shenzhen, Chinese Academy of Agricultural Sciences, Shenzhen, Guangdong 518124, China; Centre for Microbial Ecology and Genomics, Department of Biochemistry, Genetics and Microbiology, University of Pretoria, Pretoria 0028, South Africa; College of Horticulture, Academy for Advanced Interdisciplinary Studies, Nanjing Agricultural University, Nanjing, China

**Keywords:** ferns, *K*_S_ distribution, phylogenomics, gene tree – species tree reconciliation, WGD

## Abstract

Ferns, and particularly homosporous ferns, have long been assumed to have experienced recurrent whole-genome duplication (WGD) events because of their substantially large genome sizes, surprisingly high chromosome numbers, and high degrees of polyploidy among many extant members. Although, consequently, the number of sequenced fern genomes is very limited, recent studies using transcriptome data to find evidence for WGDs in ferns reached conflicting results concerning the occurrence of ancient polyploidy, for instance, in the lineage of leptosporangiate ferns. Because identifying WGDs in a phylogenetic context is the foremost step in studying the contribution of ancient polyploidy to evolution, we revisited earlier identified WGDs in leptosporangiate ferns, mainly the core leptosporangiate ferns, by building age distributions and applying substitution rate corrections and by conducting statistical gene tree – species tree reconciliation analyses. Our integrative analyses confidently identified four ancient WGDs in the sampled core leptosporangiates and suggest both false positives and false negatives for the WGDs that recent studies have reported earlier. In conclusion, we underscore the significance of substitution rate corrections and uncertainties in gene tree – species tree reconciliations in calling WGD events, and that failing to do so likely leads to incorrect conclusions.

## Introduction

Tracheophytes, or vascular plants, have shaped the diversity of the terrestrial ecosystem on Earth since their first appearance about 431 to 451 million years ago (Morris et al. 2018). Tracheophytes are composed of two major groups, spermatophytes (seed plants) and pteridophytes (non-seed vascular plants) (PPG 2016). Seed plants form a monophyletic group, including gymnosperms and angiosperms and it has been well acknowledged that paleo-polyploidizations, or ancient whole-genome duplications (WGDs), have played essential roles in their diversification and adaptation (Fox et al. 2020, Van de Peer et al. 2021). Indeed, within the past two decades strong evidence accumulated for recurrent WGDs in seed plants (Van de Peer et al. 2017) and their roles in the evolution of innovative traits and in facilitating the diversification of species (Soltis and Soltis 2016, Van de Peer et al. 2017, Landis et al. 2018). Different from the lineage of seed plants, seed-free vascular plants or pteridophytes form a paraphyletic group, including Lycopodiopsida (lycophytes) and Polypodiopsida (ferns), and for a long time, WGDs in seed-free vascular plants were indefinite, although cytological evidence suggested that polyploidization may not be uncommon in ferns (Wood et al. 2009, Clark et al. 2016, Wang et al. 2022).

Ferns are the largest group and make up more than 90% of the extant diversity of non-seed vascular plants (PPG I 2016). Compared to seed plants, very few have had their genome sequenced so far (Szövényi et al. 2021), because they, especially homosporous ferns, tend to have large genome sizes and large to huge chromosome numbers. For example, the modern fern or C-fern (*Ceratopteris richardii*) possesses n = 39 chromosomes with a genome of 11.25 Gbp (Marchant et al. 2019). More strikingly, *Ophioglossum reticulatum* is a fern species with the highest chromosome number known amongst eukaryotes, with 2n = 1,440 chromosomes (Khandelwal 1990). The huge diversity as well as the high numbers of chromosomes of ferns are compelling mysteries that have fascinated evolutionary biologists for decades (Haufler and Soltis 1986, Barker 2009, Clark et al. 2016, Wang et al. 2022). Given that polyploidizations can increase both genome size and numbers of chromosomes directly, multiple rounds of polyploidizations, along with potential changes of chromosome compositions and/or processes of genome downsizing, have been hypothesized to explain the evolution of (numbers of) chromosomes and genome sizes in ferns (Clark et al. 2016, Wang et al. 2022).

It is only until recently that genomic and transcriptomic data have begun to shed light on ancient polyploidies in ferns (Barker and Wolf 2010, Szövényi et al. 2021). Analyses of the first two genomes of heterosporous ferns, *Azolla filiculoides* and *Salvinia cucullata*, have identified two WGDs, with one specific to the genus *Azolla* and the other shared by all core Leptosporangiates (Li et al. 2018). Furthermore, the partial genome of the C-fern has also shown some evidence for a WGD in its evolutionary lineage (Marchant et al. 2019). In addition, two recent studies have added valuable supplements of transcriptome data to the scarce genomic data of ferns and suggested several ancient WGDs during the evolutionary history of ferns (1KP initiative 2019, Huang et al. 2020). However, since there have been conflicting results regarding the identified WGD events, we here revisited ancient polyploidies in ferns with state-of-the-art approaches and compared our results with those of the previous studies.

In doing so, our central focus was on the occurrence of WGDs in leptosporangiate ferns, or more specifically in the lineage of core leptosporangiates, for two reasons. First, leptosporangiate ferns form the vast majority of the species in extant ferns (Pryer et al. 2004). Leptosporangiates are subdivided into total seven orders, namely Osmundales, Hymenophyllales, Gleicheniales, Schizaeales, Salviniales, Cyatheales, and Polypodiales, the last three of which include most species and constitute the lineage of core leptosporangiates. Second, we focused on those ferns that have been investigated in previous studies with a well-resolved phylogeny (Shen et al. 2018). To this end, we selected species in all the three orders of core leptosporangiates and used representatives of another three orders from the leptosporangiates as outgroups for considering and correcting substitution rates and phylogenetic rerooting. This way, we could revisit ten out of 14 WGDs reported by the 1KP initiative (2019) and ten out of ten WGDs reported by Huang et al. (2020) in leptosporangiates. For the ten WGD events retrieved from each study, only five are congruent and have been placed in the same phylogenetic positions (Figure 1).

**Figure 1.**
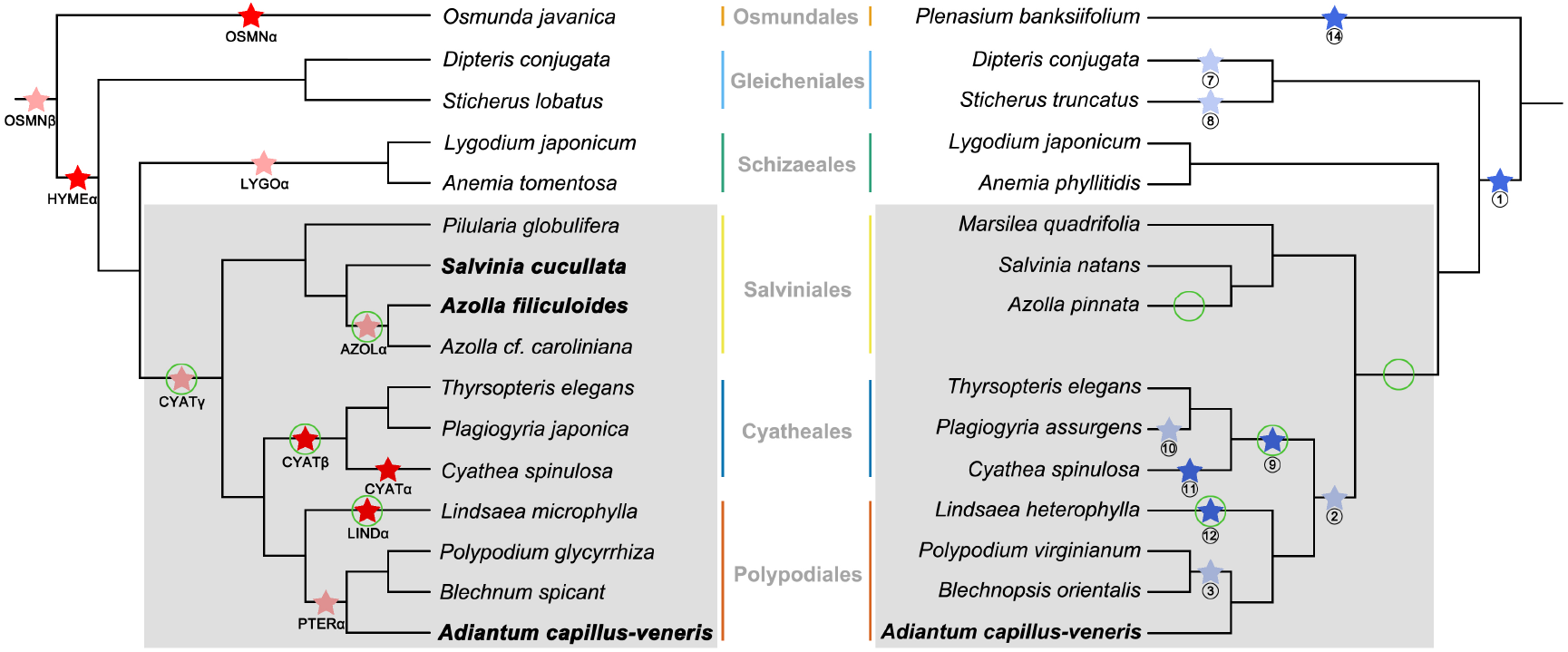
Identified WGD events in leptosporangiates, as reported by the 1KP initiative (2019) (left) and by Huang et al. (2020) (right). Ten of 14 WGDs in leptosporangiates from the 1KP initiative (2019) are denoted as (light) red stars. Four WGDs are not included because they were placed in lineages not studied by Huang et al. (2020). Ten out of ten WGD events from Huang et al. (2020) are denoted as (light) blue stars. WGD events found in the same lineages by both studies are in solid red and solid blue. The grey background highlights the core leptosporangiates in the two phylogenetic trees. The names for species with fully sequenced genomes are in bold. The green circles denote the WGDs in core leptosporangiates identified in this study.

The conflicting results with respect to the identified WGD events in previous studies could result from several pitfalls that are often overlooked in the commonly used approaches to find evidence for ancient WGDs. Indeed, both studies previously mentioned utilized the construction of paralogous age distributions and gene tree – species tree reconciliation. Although these two approaches have great power and have been widely applied to detect WGDs based on genomic and transcriptome data (Jiao et al. 2011, Vanneste et al. 2013, Li et al. 2015, McKain et al. 2016, Zhang et al. 2017, Ren et al. 2018), they must be used with caution (Tiley et al. 2018, Zwaenepoel et al. 2019, Zwaenepoel and Van de Peer 2019). WGDs in so-called *K*_S_-age distributions, where the number of duplicates is plotted against their age (as inferred from the expected number of synonymous substitutions per synonymous site, i.e., the synonymous distance *K*_S_ (Vanneste et al. 2013)), can be identified with peaks in the distribution, which suggest that many genes have been duplicated at the same time in evolutionary history. Such *K*_S_ peaks are often compared with speciation events characterized by *K*_S_ distributions based on orthologs between species to infer the relative or absolute timing of the large-scale duplication event. However, such comparisons admit meaningful interpretation only if substitution rates of the species under consideration are similar, while substitution rates naturally vary among lineages. It has been gradually acknowledged that different substitution rates can affect the placement of WGDs through comparisons between WGD events and speciation events (Barker et al. 2008; Chen et al. 2020, Sensalari et al. 2021). For instance, if two species diverged and have evolved at different substitution rates after their common ancestor shared a WGD event and the difference in substitution rates is not accounted for, the WGD *K*_S_ peak identified in the species with a lower substitution rate may be incorrectly interpreted as a younger and lineage-specific WGD. In contrast, species with a higher substitution rate may still support a shared WGD. This could eventually lead to erroneous conclusions, especially when no genome is available to determine the inference of WGDs via collinear or synteny analysis.

A second approach to identify and date WGDs is to use gene tree – species tree reconciliation, where events underlying the evolutionary history of a gene, like gene duplication and loss, hybridization, introgression, horizontal gene transfer, and incomplete lineage sorting, are identified by mapping gene trees onto species trees. When many duplicated genes are reconciled on one specific branch of the species tree, this can be considered evidence for a large-scale or WGD event. Although there are differences in the way gene tree – species tree reconciliation approaches have been implemented in the previously mentioned studies (1KP initiative (2019), Huang et al. (2020)), both have employed the least common ancestor (LCA) reconciliation to determine duplication events on a species phylogeny based on gene trees inferred using maximum likelihood (ML) inference. In LCA reconciliation, a duplication event involving genes from some species is placed on a species phylogeny at (we say ‘reconciled to’) the node associated with the most recent common ancestor of these species (Zmasek and Eddy, 2001). Even if gene trees have been filtered based on their quality before reconciliations (using some criterion based on bootstrap support values for instance), the LCA reconciliation is still error-prone, and its accuracy depends on the correctness of inferred gene tree topologies (Hahn 2007). Nevertheless, the true gene tree topology for a gene family is often one among many statistically equivalent gene trees (Wu et al. 2013), so only considering the one ‘best’ ML tree for each gene family may cause systematic bias when using LCA reconciliation to identify WGDs (Hahn 2007; Zwaenepoel and Van de Peer 2019). In addition, a WGD event and its phylogenetic position are often determined when the number of duplication events on a branch exceeds a certain cut-off, which is usually set somewhat arbitrarily without acknowledging the varying contribution of small-scale duplications (SSDs) along different branches of the species tree (Li et al. 2015, McKain et al. 2016, Ren et al. 2018), which may result in false positive WGD identification towards the tips of a species phylogeny (Zwaenepoel and Van de Peer 2019).

To revisit ancient polyploidy in leptosporangiate ferns, we retrieved relevant transcriptome data from the 1KP initiative (2019), for its relatively high quality and reasonable gene numbers (Supplementary Figure 1). Also, we added two publicly available genomes of heterosporous ferns and a newly sequenced homosporous fern, *Adiantum capillus-veneris* L. (Fang et al., 2022, accepted pending revisions). We did not include the genome of the C-fern because of its partial and fragmented nature (Marchant et al. 2019). By considering differences in substitution rates and performing statistical gene tree – species tree reconciliations under a model integrating both gene duplication and loss (DL) and large-scale gene duplication (WGD) and loss, we confidently identified four WGDs in core leptosporangiate ferns (Figure 1), fewer than the six WGDs as found by 1KP initiative (2019) and Huang et al. (2020), while some WGDs have also been predicted at different phylogenetic positions, suggesting that some WGDs identified by the two previous studies are likely false positives. Our study again highlights the importance of fully recognizing the caveats and limitations of commonly used approaches in calling WGD events.

## Results and Discussion

### Different substitution rates among ferns in core leptospongiates

To compare synonymous substitution rates among core leptosporangiate ferns, we compared one-to-one orthologous *K*_S_ distributions between *Lygodium japonicum* from Schizaeales and species from Cyatheales, Salviniales, and Polypodiales, the three orders within core leptospongiates (Figure 2a–c). Because peaks in the orthologous *K*_S_ distributions all represent the same event, i.e., the divergence between Schizaeales and the core leptospongiates, they should have identical or at least very similar *K*_S_ values if the selected species all have similar substitution rates. However, our results show that the orthologous *K*_S_ peak values are smaller for species from Cyatheales than species from Salviniales and Polypodiales, suggesting that Cyathealean species have slower substitution rates than species of the other two orders in the core leptosporangiates. In addition, the orthologous *K*_S_ peak values for the species belonging to Salviniales and Polypodiales show more variation than the ones in Cyatheales, suggesting that they tend to have more variable substitution rates between the different species (Figure 2a–c). We also found similar patterns of differences in synonymous substitution rates among the three orders in core leptosporangiates, when using *Dipteris conjugata* from Gleicheniales as an outgroup for the analyses (Supplementary Figure 2).

**Figure 2.**
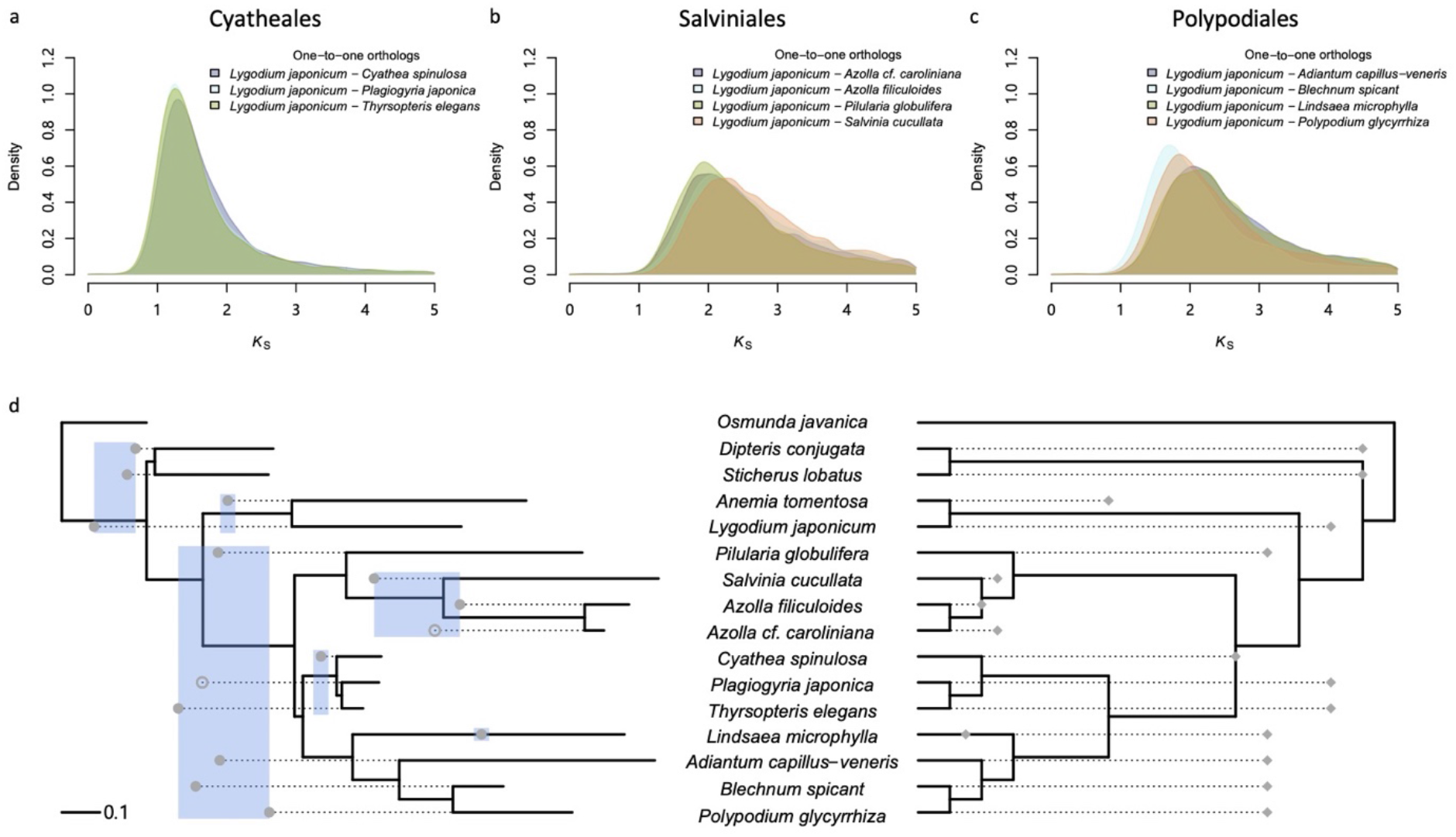
Orthologous age distributions and WGD events identified based on *K*_S_ distributions for the whole paranomes of different fern species. (a-c) One-to-one orthologous *K*_S_ distributions between *L. japonicum* and species from Cyatheales (a), Salviniales (b), and Polypodiales (c); (d) WGD events identified based on *K*_S_ distributions for the whole paranome of species in the phylogeny. On the left, a species phylogram is shown with branch lengths in *K*_S_ units, while WGD events are depicted as dots (calculated as half the *K*_S_ peak value of each species starting from the corresponding tip). Solid dots denote significant *K*_S_ peaks in the SiZer analysis, whereas hollow dots denote a *K*_S_ peak only identified by GMM but not SiZer (see Methods and Supplementary Figures 3 and 4). On the right, a species cladogram is shown, where WGD events are depicted as rhombs according to the analyses of ksrates (Sensalari et al. 2021). Note that when a WGD and a speciation event overlap in the ksrates analysis, the WGD event is placed at the speciation event in the cladogram.

As substitution rates affect *K*_S_ distributions, peaks in the paralogous *K*_S_ distributions may differ for species with different substitution rates, even if they have experienced the same WGD event. We hence wondered if the differences in calling WGDs in the core leptosporangiates in previous studies could be due to different substitution rates. To this end, we first built *K*_S_ distributions for the full paranomes (i.e., all the paralogous genes) of 16 selected species (Supplementary Figures 3 and 4). Peaks in the *K*_S_ distributions, which potentially represent WGDs, were identified by Gaussian Mixture Modelling (GMM) and verified by the SiZer analyses to distinguish significant peaks from non-significant ones (see Methods). In all species, except for *Plagiogyria japonica* and *Azolla cf. caroliniana*, GMM discovered a peak supported by the SiZer analyses (Supplementary Figure 3). The peaks show various *K*_S_ values and are largely in line with the paranome *K*_S_ distributions from the 1KP initiative (2019).

To correct for different synonymous substitution rates among species, we adopted two recently developed approaches. In the first approach, the synonymous substitution rate of a species is ‘adjusted’ based on a relative rate comparison to its sister lineage in the phylogenetic tree (Figure 2d). This way, we can adjust a WGD peak starting from the corresponding tip in the species tree (Chen et al. 2020). To implement this, we first inferred a phylogenetic tree based on 34 single-copy genes identified from the 16 species, with branch lengths in *K*_S_ units (see Methods). If we assume that both paralogs, on average, evolved at a similar rate after a WGD event, we could simply consider half the *K*_S_ values of all identified peaks in each species to position, starting from each tip, the WGDs on the phylogenetic tree (Figure 2d). In the second approach, we used ksrates, which adjusts synonymous substitution rates to the rate of a focal species by relative rate tests (Sensalari et al. 2021). Therefore, peaks identified in the *K*_S_ age distribution of the focal species can be directly compared with speciation events represented by *K*_S_ distributions based on orthologs (Supplementary Figure 5).

By applying the first approach, our results show that considering various synonymous substitution rates among lineages is essential in correctly interpreting the identified *K*_S_ peaks. For instance, both the 1KP initiative (2019) and Huang et al. (2020) identified two WGDs within Polypodiales. One WGD is placed in the lineage of *Lindsaea* and is supported by both studies (‘LIND𝛼’ and ‘12’ in Figure 1), as well as by our *K*_S_ analyses (Figure 2d). The 1KP initiative (2019) suggests another WGD (‘PTER𝛼’ in Figure 1) shared by *Polypodium*, *Blechnum*, and *Adiantum*, whereas Huang et al. (2020) suggests a WGD (‘3’ in Figure 1) only shared by *Polypodium* and *Blechnum* (Figure 1). However, in our results, considering differences in substitution rates, the *K*_S_ peaks found in *Blechnum spicant*, *Polypodium glycyrrhiza*, and *Adiantum capillus-veneris,* all support a more ancient WGD likely shared by the core leptospongiates (Figure 2d), although they all have different *K*_S_ peak values (Supplementary Figure 3) because *Blechnum spicant* and *Polypodium glycyrrhiza* have lower substitution rates than *Adiantum capillus-veneris* and *Lindsaea microphylla* (Figure 2c; Supplementary Figure 2).

In Cyatheales, both the 1KP initiative (2019) and Huang et al. (2020) support a WGD in *Cyathea* (‘CYAT𝛼’ and ‘11’ in Figure 1) and a shared WGD for Cyatheales (‘CYATβ’ and ‘9’ in Figure 1). In addition, according to Huang et al. (2020), there is another WGD in the lineage leading to *Plagiogyria* (‘10’ in Figure 1). However, the *K*_S_ peaks identified in our analyses neither support a WGD in *Cyathea* (‘CYAT𝛼’ and ‘11’ in Figure 1) or in *Plagiogyria* (‘10’). Instead, the *K*_S_ peak we found in *Cyathea spinulosa* supports a WGD shared by Cyathales, and the peak we found in *Thyrsopteris elegans* even supports a WGD before the divergence between the core leptospongiates and Schizaeales (Figure 2d). Although the paranome *K*_S_ distribution of *Plagiogyria japonica* shows no significant peak according to the SiZer analysis, the GMM still shows a less perceptible peak suggesting an ancient WGD event in *Thyrsopteris elegans*. Evidently, the three species in Cyatheales have the lowest substitution rates among the core leptosporangiates (Figure 2a). Therefore, both the 1KP initiative (2019) and Huang et al. (2020) might have misinterpreted the peak in *Cyathea spinulosa* at *K*_S_ ≈ 0.3 as evidence for a recent WGD event in the genus of *Cyathea*, while this peak is actually the result of a more ancient WGD, with the small *K*_S_ values due to the comparatively low substitution rates. If there has been a WGD shared by Cyatheales, we would expect to observe clear *K*_S_ peaks in *Thyrsopteris elegans* and *Plagiogyria japonica* as well. Unexpectedly, only a *K*_S_ peak in support of an even more ancient WGD has been observed in *Thyrsopteris elegans*. We argue that *Thyrsopteris elegans* and *Plagiogyria japonica* have even lower synonymous substitution rates than does *Cyathea spinulosa* (Figure 2a). If the *K*_S_ peak for the Cyathealean WGD is at *K*_S_ ≈ 0.3 in the paranome *K*_S_ distribution of *Cyathea spinulosa*, the expected *K*_S_ peaks in slower evolving *Thyrsopteris elegans* and *Plagiogyria japonica* must then have even smaller values, which may be confounded by the background *K*_S_ distribution from SSDs.

In Salviniales, Huang et al. (2020) found no evidence for WGD, but the 1KP initiative (2019) identified one WGD in the lineage leading to *Azolla*, which is consistent with the observation made in the paper presenting the genome sequences of *Salvinia cucullata* and *Azolla filiculoides* (Li et al. 2018). Similarly, we found a peak in the paranome *K*_S_ distribution of *Azolla filiculoides* that supports a WGD. However, the paranome *K*_S_ distribution of *Salvinia cucullata* has a peak supporting a WGD before the divergence of *Salvinia* and *Azolla,* instead of a WGD before the divergence of core leptosporangiates, as suggested earlier (Li et al. 2018). Also, the GMM has disentangled a peak in the paranome *K*_S_ distribution of *Azolla cf. caroliniana* suggesting a WGD shared by *Salvinia* and *Azolla*, but the peak is not significant in the SiZer analysis (Supplementary Figure 3). Although the above results indicate two competing phylogenetic placements of WGDs within Salviniales, it is clear that a WGD shared by core leptosporangiates is evidenced by the *K*_S_ peak found in *Piluaria globulifera*. Note that in each of the three orders of core leptosporangiates, there is at least one species that lends support for an ancient WGD shared by all the core leptosporangiates (or even before the divergence between Schizaeales and core leptosporangiates) as identified by the 1KP initiative (2019) (‘CYAT𝛾’ in Figure 1). In contrast, Huang et al. (2020) has indicated a WGD shared by Polypodiales and Cyatheales (‘2’ in Figure 1), which has no support from the 1KP initiative (2019), nor from our results.

Additionally, outside the lineage of core leptosporangiates, the *K*_S_ peak identified in *Anemia tomentosa* indicates a shared WGD with *Lygodium japonicum*. However, the *K*_S_ peak in *Lygodium japonicum* goes against a shared WGD with *Anemia tomentosa* but suggests a more ancient WGD, which is also supported by the *K*_S_ peaks in *Dipteris conjugate* and *Sticherus lobatus* (Figure 2d). In general, for these species our results seem to largely agree with the 1KP initiative (2019), which identified a WGD in *Anemia tomentosa* (‘LYGO𝛼’ in Figure 1), but no WGDs in *Dipteris* and *Sticherus*, in contrast to those in Huang et al. (2020) (‘7’ and ‘8’ in Figure 1). For the more ancient WGDs identified by both previous studies (‘OSMNβ’, ‘HYME𝛼’, and ‘1’ in Figure 1), our analyses could not resolve whether a WGD occurred before the divergence of leptosporangiates (‘OSMNβ’ in Figure 1) and/or a WGD occurred after *Osmunda javanica* diverged from the rest of leptosporangiates (‘HYME𝛼’ and ‘1’ in Figure 1). The reason we could not resolve this is because *Osmunda javanica* is an outgroup in the phylogenetic tree (in *K*_S_ units) and without extra information we cannot determine when *Osmunda javanica* diverged from other leptosporangiates. This is also why, although there is a *K*_S_ peak in the distribution for *Osmunda javanica* (Supplementary Figure 3), we were uncertain about assuming a WGD either shared with other leptosporangiates or a species-specific WGD (‘OSMN𝛼’ and ‘14’ in Figure 1). Although we could add extra species to determine the root on the branch leading to *Osmunda javanica*, this may introduce another species with a WGD event that cannot be resolved with certainty, so here we decided to focus our analyses on the core leptosporangiates.

By applying the second approach, using ksrates, to correct for unequal substitution rates among species (Sensalari et al. 2021), we can directly compare the WGD peaks identified in the paranome *K*_S_ distribution of a focal species with speciation events represented by peaks in *K*_S_ distributions of orthologs between species (Supplementary Figure 5), and hence place the identified WGDs on a cladogram of the species tree (Figure 2d). Our previous results are confirmed, except that the ksrates analysis provides extra support from *Lindsaea microphylla* for a WGD shared by the core leptospongiates. In summary, both above analyses of *K*_S_ distributions, considering substitution rate corrections, show that different substitution rates among lineages affect the placement of WGDs in a phylogenetic context. Not considering substitution rate differences may therefore result in false positives or incorrectly called WGDs.

### Evaluating WGDs using a small- and large-scale gene duplication – gene loss model

Although analyses based on *K*_S_-age distributions and considering different substitution rates in different species could already reject some of the WGDs proposed in earlier studies, some *K*_S_ peaks for different species fall in competing branches adjacent to each other in the species tree (Figure 2d) and remain therefore ambiguous. For example, the *K*_S_ peaks from *P. glycyrrhiza*, *P. globulifera*, and *A. capillus-veneris* support a WGD shared by all the core leptospongiates, whereas the ones from *T. elegans*, *B. spicant*, and *P. japonica* support a WGD before the divergence between the core leptospongiates but also Schizaeales. Similarly, the *K*_S_ peak from *Azolla filiculoides* supports a WGD specific to *Azolla*. Still, the *K*_S_ peak for *Salvinia cucullata* favors a shared WGD by *Azolla* and *Salvinia*, the two heterosporous fern genera. These results could point to two independent WGDs, one before and one after the speciation event, or alternatively, to one WGD event that is however represented by *K*_S_ peaks with different *K*_S_ peak values for different species. Specifically, for species with high substitution rates (Figure 2a–c), identifying peaks representing an ancient WGD at a large *K*_S_ value may be confounded by substitution saturation to different extents (Vanneste et al. 2013), leading to less accurate estimates of *K*_S_ peak values consequently.

Besides the uncertainty of WGD events discussed in the previous paragraph, *K*_S_-age distributions show evidence for WGDs on another four branches in the species phylogeny (Figure 3). To try to determine the exact position of all these potential WGDs, we used the so-called DL+WGD model implemented in Whale which considers gene duplication and loss due to both small-scale duplications (SSDs) and WGD (Zwaenepoel and Van de Peer 2019). Different from WGDgc, using gene counts in the DL+WGD model (Rabier et al. 2013), Whale performs probabilistic gene tree – species tree reconciliation using amalgamated likelihood estimation (Szöllõsi et al. 2013) to test WGD hypotheses in a phylogenetic context through estimating duplication (*λ*) and loss (*μ*) rates for SSDs, and duplicate retention rates (*q*) of WGDs (Zwaenepoel and Van de Peer 2019). Because Whale can consider gene tree uncertainty in its statistical reconciliation algorithm, we built gene trees for 6,863 gene families with MrBayes (Ronquist et al. 2012) and fed Whale with 10,000 trees sampled every ten generations over a total of 100,000 trees sampled from the posterior for each gene family (see Methods). To set up the DL+WGD model in Whale (see Methods for the choice of prior distributions), the eight hypothetical WGDs according to the *K*_S_ analyses were placed on the species tree (Figure 3 and Supplementary Figure 7), each with a uniform prior for the WGD retention rate. Because assuming constant duplication and loss rates of SSDs across the species tree could substantially affect WGD testing (Zwaenepoel and Van de Peer 2019), we adopted two models to incorporate various DL rates of SSDs across the species tree: 1) the critical branch-specific model, where each branch in the species tree has an equal rate for duplication and loss, i.e., *λ* = *μ* (called the ‘turnover’ rate in e.g. CAFE (De Bie et al. 2006)), but the rates vary across branches (we assume a relaxed DL clock); and 2) the relaxed branch-specific model, where duplication and loss rates again vary across branches and now are not necessarily equal, i.e., *λ* ≠ *μ*. The reason for adopting two different models is that the basic linear birth-death process DL model may not be an ideal model of gene family evolution (Zwaenepoel and Van de Peer 2021), so that comparing results from different models may aid in assessing the robustness of particular inferences to model violations. Finally, because most of the analyzed species only have transcriptome data, we used the missing values in the BUSCO analyses of each species (Supplementary Figure 6) as a parameter for taking missing genes into account in the models.

**Figure 3.**
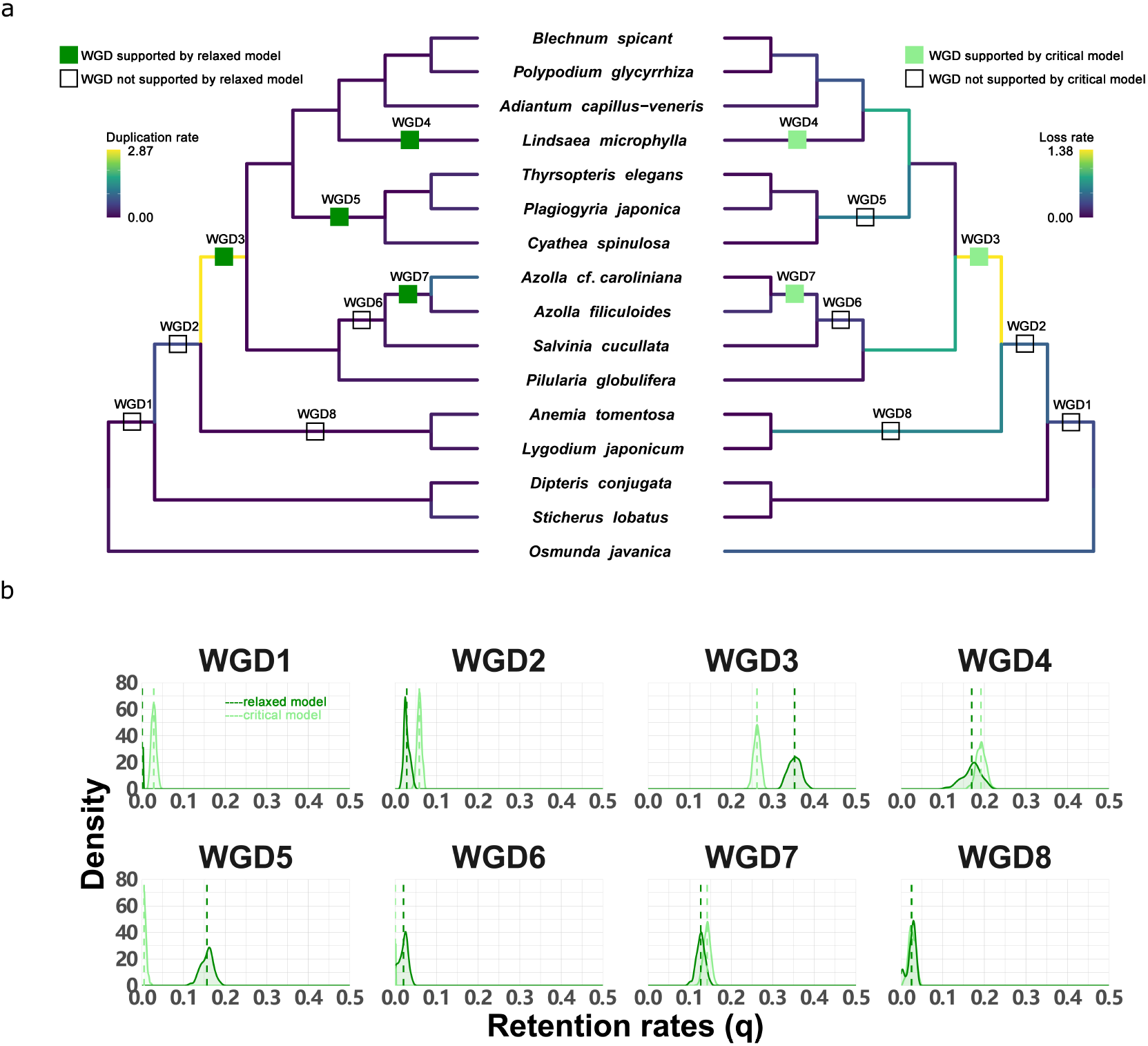
Whale (gene tree – species tree reconciliation) analysis for eight hypothetical WGDs under the DL+WGD model. (a) The species cladograms with the eight putative WGD events mentioned in the previous *K*_S_-age analyses. The WGD bars in green on the left cladogram (for the relaxed branch-specific model) and in light green on the right cladogram (for the critical branch-specific model) are supported WGDs with retention rates significantly different from zero, while the hollow WGD bars in each cladogram are the ones with retention rates not different from zero (Table 1). Posterior mean of duplication (left) and loss (right) rates estimated under the relaxed DL+WGD model (see Methods) are colored on the cladograms. Panel (b) shows the posterior distributions of the WGD retention rates (*q*) for the eight putative WGDs. The dotted lines show the posterior mean of each posterior distribution. Color code is the same as the WGD bars in the phylogenies in (a).

**Table 1.** Hypothetical WGDs, posterior mean of duplicate retention rate (*q*), and the Bayes Factor (*K*) to compare the likelihood of *q* = 0 (H_0_) to the likelihood of *q* > 0 (H_1_) using the Savage-Dickey density ratio.

| Hypotheses | Relaxed branch-specific model |  | Critical branch-specific model |  |
| --- | --- | --- | --- | --- |
| | $\bar{q}$ | $K$ | $\bar{q}$ | $K$ |
| WGD1 | 0.001 | 1272.297 | 0.029 | 2.693 |
| WGD2 | 0.028 | 2.649 | 0.058 | 0.393 |
| WGD3 | 0.352 | 0.047** | 0.263 | 0.061** |
| WGD4 | 0.170 | 0.197* | 0.193 | 0.094** |
| WGD5 | 0.156 | 0.149* | 0.006 | 128.480 |
| WGD6 | 0.020 | 21.026 | 0.000 | 6919.539 |
| WGD7 | 0.127 | 0.166* | 0.142 | 0.135* |
| WGD8 | 0.025 | 12.864 | 0.027 | 6.007 |

\*\* $K < 1/10$ or $K < 0.1$ , strong evidence against $H_0$ ;
\* $K < 1/10^{0.5}$ or $K < 0.3162$ substantial evidence against $H_0$ ;
$K < 1$ , $H_1$ supported, not worth more than a bare mention; $K > 1$ , $H_0$ supported.

After obtaining posterior distributions of all the parameters under both DL+WGD models (Figure 3), we estimated the duplicate retention rate (*q*) of each putative WGD by its posterior mean. Unlike using the DL+WGD model in an ML scheme (e.g., WGDgc), in which a series of likelihood ratio tests were performed by removing only one WGD at a time to test its likelihood of occurrence (Tiley et al. 2016), we used the posterior distributions of *q* to estimate the Bayes Factor (*K*) to test if *q* is significantly different from zero using the Savage-Dickey density ratio (Zwaenepoel and Van de Peer 2019). A putative WGD with an estimated value of *q* significantly larger than zero would hence indicate the occurrence of a WGD on a specific branch (Table 1). With the relaxed DL+WGD model, our results support four WGDs, i.e., WGD3, WGD4, WGD5, and WGD7, which all have a WGD retention rate over 0.05. Similarly, the results based on the critical branch-specific DL+WGD model support WGD3, WGD4, and WGD7 (Figure 3). Besides, WGD1 and WGD8 obtained no support from Whale and they were placed in the outgroups of our focal leptosporangiate ferns, so the species sampling may be less suitable to resolve these WGDs as discussed above. For example, WGD1 may be the result of two WGDs that have occurred on two consecutive branches if we accept the results from the 1KP initiative (2019). Without species that can further break down the branch in the species phylogeny where WGD1 was located, it is difficult to neatly solve the problem with either the *K*_S_ or the reconciliation approaches (Zwaenepoel and Van de Peer 2019).

Our gene tree – species tree reconciliation analyses with both DL+WGD models raised our confidence in resolving the two WGDs discussed higher, i.e., WGD6 or WGD7, and WGD2 or WGD3. Also, the support of a WGD in the lineage leading to *Lindsaea* (WGD4) is decisive, while the WGD shared by Cyatheales (WGD5) requires further discussion on the performance of the critical and relaxed branch-specific models. Below, we discuss the four WGDs in core leptosporangiates in more detail.

### *The* Azolla *WGD*

Both the relaxed and critical branch-specific models strongly support a WGD in the lineage leading to *Azolla* (WGD7) rather than a WGD shared by *Azolla* and *Salvinia* (WGD6) (Figure 3 and Table 1). Although the latter seems to have some support from the *K*_S_ analyses (Figure 2d), the peak in *Azolla cf. caroliniana* did not show to be significant in the SiZer analysis (Supplementary Figure 3), and the peak in *S. cucullata* may be artificial due to substitution saturation, because it has the highest substitution rate among the analyzed species in Salviniales (Figure 2b). Because the genomes of *A. filiculoides* and *S. cucullata* are available (Li et al. 2018), we studied intra-genomic collinearity in each species and identified paralogous genes located in collinear regions. In addition to analyzing the *K*_S_ distributions for anchor pairs (pairs of genes still residing in collinear regions of the genome; Supplementary Figures 8 and 9), we examined the Whale reconciliation results for anchor pairs (see Methods). Note that the reconciliation result of a pair of paralogs in Whale is not a duplication event on a specific branch but a posterior distribution over the possible branches in the species phylogeny where the duplicate may reconcile to (Figure 4). Except for some anchor pairs reconciled with high posterior probability to the species-specific branch, anchor pairs from the *A. filiculoides* genome tend to support the WGD specific to the *Azolla* genus (WGD7) rather than a WGD shared by *Azolla* and *Salvinia* (WGD6) (Figure 4a). Also, the reconciliation results for anchor pairs from *S. cucullata* only lend little support for a WGD shared by *Azolla* and *Salvinia* (Figure 4b). In addition, a further inter-genomic collinearity comparison shows that the syntenic ratio of *A. filiculoides*: *S. cucullata*: *A. capillus-veneris* is 2: 1: 1, which again confirms our conclusion of one round of WGD experienced by *Azolla,* while no evidence for a WGD on the branch leading to *Adiantum* and *Salvinia* (see Methods and Supplementary Figure 12).

**Figure 4.**
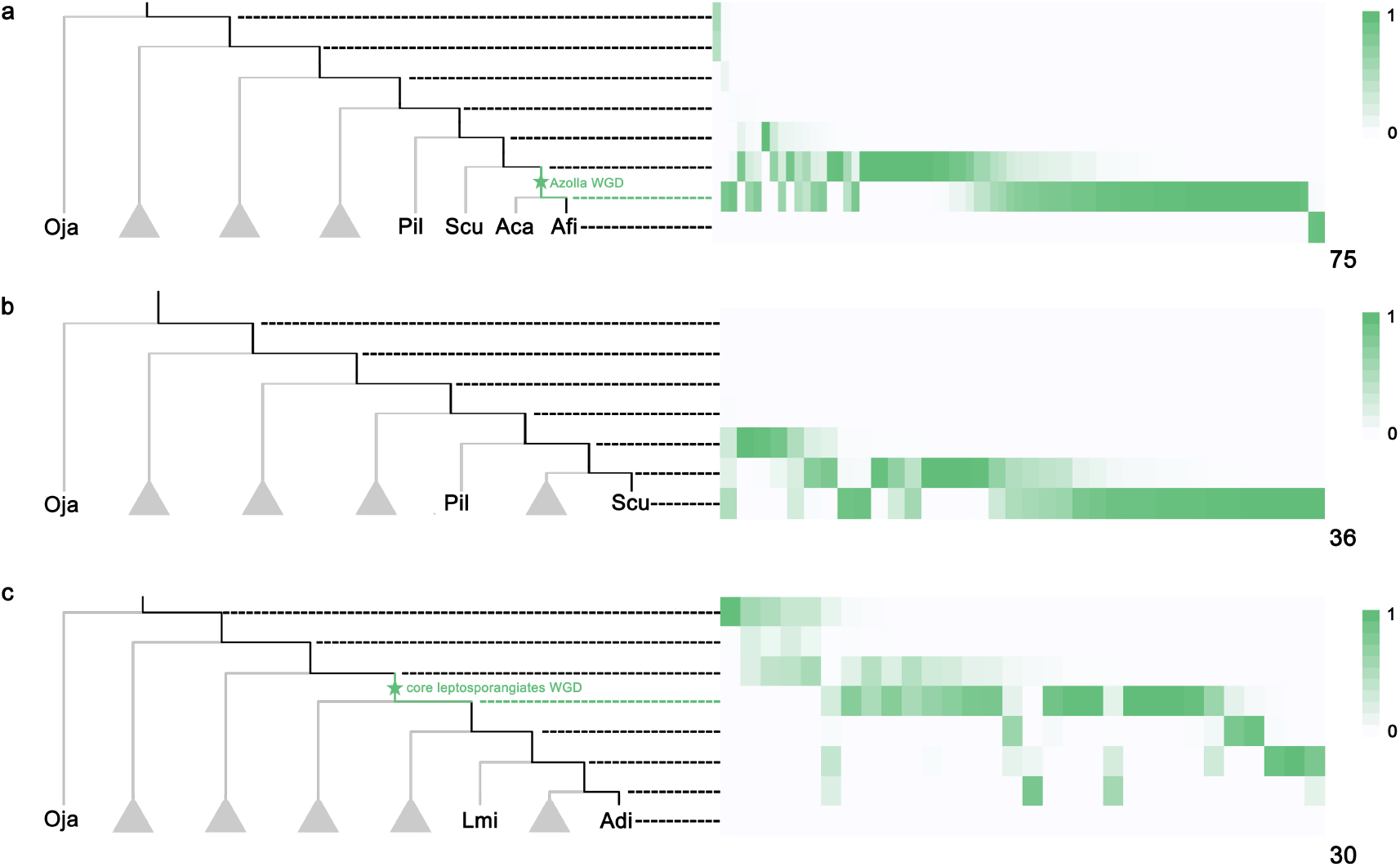
Gene tree – species tree reconciliation analyses for anchor pairs identified in the three fern genomes, *A. filiculoides*, *S. cucullata*, and *A. capillus-veneris*. On the phylogenetic trees, branches highlighted in black are the ones to which an anchor pair in the genomes of (a) *A. filiculoides*, (b) *S. cucullata*, and (c) *A. capillus-veneris* can be reconciled in the gene tree – species tree reconciliation analyses. On the right, the total number of columns in each heatmap, denoted at the right bottom corner, is the number of anchor pairs in analyzed (see Methods). The squares in white to green in each column show the posterior probability that an anchor pair is reconciled as a duplication event to the respective branch. The color code ranges from white (posterior probability equal to zero) to green (posterior probability equal to one).

### The WGD shared by core leptosporangiates

With respect to WGD2 and WGD3, our Whale results support the WGD shared by core leptosporangiates (WGD3) but reject a WGD before the divergence between core leptosporangiates and Schizaeles (WGD2) (Figure 3 and Table 1). In the critical branch-specific model, the posterior mean for WGD2 is over 0.05 and larger than that in the relaxed branch-specific model. Also, although the Bayes Factor of WGD2 slightly favors a duplicate retention rate over zero, it cannot provide strong evidence for the occurrence of a WGD (Table 1). By further examining the anchor pairs in the *A. capillus-veneris* genome (Supplementary Figure 10), we found that the Whale reconciliation results for the anchor pairs also only support a WGD shared by the core leptosporangiates (Figure 4c).

### *One WGD in Cyatheales and one WGD in* Lindsaea

The support of a WGD shared by Cyatheales is not as decisive as the one in the lineage leading to *Lindsaea*. The latter is supported by both DL+WGD models, as well as by the *K*_S_ analysis of *Lindsaea microphylla*. However, the former is only supported by the *K*_S_ peak in *Cyathea spinulosa*, but not in the other two species, *T. elegans* and *P. japonica*. In addition, the critical branch-specific model has an estimate for a low WGD retention rate ≈ 0, compared with the WGD retention rate of 0.16 in the relaxed branch-specific model. In the relaxed model, the duplication rate is low 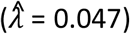, but the loss rate is high 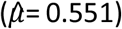 on the branch leading to Cyatheales, so in the critical branch-specific model, the duplication rate is higher, whereas the loss rate is lower compared to the two rates estimated by the relaxed branch-specific model, respectively 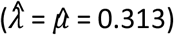 (Supplementary Figure 7). Therefore, on the branch with WGD5 (Figure 3), the assumption of equal duplication and loss rates in the critical branch-specific model appears to be strongly violated, although at the same time the rate differences in the branch-specific model appear to be unrealistic when interpreted as a model of gene family evolution. It seems prudent to conclude that support for WGD5 is not robust to model violations, and to abstain from further judgment on the basis of these phylotranscriptomic analyses. These issues highlight the problems with phylogenomic modeling of gene family evolution, and the need for more realistic models for genome evolutionary processes.

## Conclusions

Accurate identification of WGDs in a phylogenetic context is the first and vital step to studying the consequences of ancient polyploidy during evolution. Lacking high-quality genome assemblies, identifying WGDs in seed-free vascular plants has been primarily based on paranome age distributions and gene tree – species tree reconciliations using transcriptome data. By revisiting both genomic and transcriptome data for leptosporangiate ferns, especially core leptosporangiates, we showed that neglecting differences in substitution rates and performing LCA reconciliations could lead to false positives and false negatives in calling WGDs. Therefore, we underscore the importance of careful analysis, including the consideration of differences in substitution rates and appreciation of gene tree – species tree reconciliation uncertainties, prompting that failure to do so is likely to lead to unreliable or incorrect conclusions. In addition, we highlight the importance of developing better and more robust statistical models for genome evolutionary processes if we are to reliably characterize the evolutionary history of life on earth at the genomic level.

## Acknowledgements

YVdP acknowledges funding from the European Research Council (ERC) under the European Union’s Horizon 2020 research and innovation program (No. 833522) and from Ghent University (Methusalem funding, BOF.MET.2021.0005.01). HC acknowledges funding from the Research Foundation – Flanders (FWO) (No. 3G032219). AZ acknowledges the PhD Fellowship of FWO.

## Materials and Methods

### Transcriptomes and Genomes of Leptosporangiates

To revisit WGDs identified in previous studies, we selected 16 and 15 species and their corresponding assembled transcriptomes from the 1KP initiative (2019) and Huang et al (2020), respectively. Except for the order Hymenophyllales (due to its uncertain phylogenetic position (1KP initiative 2019, PPG I 2016)), the sampled species were from the remaining six orders in Polypodiidae (leptosporangiate fern), including Osmundales, Gleicheniales, Schizaeales, Salviniales, Cyatheales, and Polypodiales (Figure 1 and Supplementary Table 1). Unigenes with identical sequences were removed by SeqKit (Shen et al. 2016) to eliminate redundancy. Because of the relatively high quality and reasonable gene numbers (Supplementary Figure 1), we only used the transcriptome data from the 1KP initiative (2019) for further analyses. The genomes of *Azolla filiculoides* and *Salvinia cucullata* were retrieved from fernbase.org (Li et al. 2018), and the genome of *Adiantum capillus-veneris* was obtained from our collaborators (Fang et al., 2022, accepted pending revision). Only the longest transcripts were retained for genes from the three genomes if more than one isoform was present at a gene locus. All coding sequences of genes were further filtered to assure that they were divisible by three and had no unknown nucleotides or premature stop codons. We then used BUSCO (Simão et al. 2015) to assess the completeness of gene space in the 16 selected ferns utilizing the database of embryophyta_odb10 (Kriventseva et al. 2019) (Supplementary Figure 6).

### Constructing K_S_-based age distributions

*K*_S_-based age distributions for all paralogous genes (paranome) in transcriptomes and genomes were constructed by wgd (Zwaenepoel and Van de Peer 2018). In brief, the paranome was built by identifying gene families with the mclblastline pipeline (v.10-201) (Van Dongen 2000) with an inflation factor of 2.0 after performing all-against-all BLASTP (v.2.6.0+) (Camacho et al. 2009) search with an *E*-value cut-off of 1 × 10^−10^. Each gene family was aligned using MUSCLE (v.3.8.31) (Edgar 2004) and CODEML in the PAML package (v.4.9j) (Yang 2007) was used to estimate *K*_S_ for all pairwise comparisons within a gene family. As a gene family of n members produces n(n − 1)/2 pairwise *K*_S_ estimates for n − 1 retained duplication event, wgd corrected for the redundancy of *K*_S_ values by first inferring a phylogenetic tree for each family using Fasttree (v.2.1.7) with the default settings (Price et al. 2010). Then, for each duplication node in the resulting phylogenetic tree, all m *K*_S_ estimates for a duplication between the two child clades were added to the *K*_S_ distribution with a weight of 1/m, so that the sum of the weights of all *K*_S_ estimates for a single duplication event was 1.

To detect peaks in the *K*_S_ distributions that could be signatures of WGD events, we performed mixture modeling using the R package mclust (Scrucca et al. 2016). We first transformed *K*_S_ distributions into log-scale, which were further fitted to mixture models of Gaussian distributions (Rasmussen 1999). We increasingly fitted one to eight components per mixture model and used the Bayesian Information Criterion (BIC) to select the optimal number of components. Although BIC strongly penalizes increases in the number of model parameters, the Gaussian mixture modeling is still prone to overfitting, so we further performed SiZer (Significance of Zero Crossings of the Derivative) analysis using the R package feature (https://cran.r-project.org/web/packages/feature/index.html) to distinguish *bona fide* peaks in the *K*_S_ distributions from those that represent noises (Chaudhuri and Marron 1999) (Supplementary Figure 3).

### Correcting differences in synonymous substitution rates

The *K*_S_-based orthologous age distributions were constructed by wgd (Zwaenepoel and Van de Peer 2018), which identified one-to-one orthologs followed by *K*_S_ estimation using the CODEML program, as described above. To compare different synonymous substitution rates in core leptosporangiate species, we compared the *K*_S_ distributions of one-to-one orthologs identified between *Lygodium japonicum* and *Cyathea spinulosa*, *Plagiogyria japonica*, and *Thyrsopteris elegans* from Cyatheales, *Salvinia cucullata*, *Azolla cf. caroliniana*, *Azolla filiculoides*, and *Pilularia globulifer* from Salviniales, *Adiantum capillus-veneris*, *Blechnum spicant*, *Lindsaea microphylla*, and *Polypoddium glycyrrhiza* from Polypodiales (Figure 2a–c). Because *L. japonicum* and the above core leptosporangiate ferns diverged at a specific time, we would expect similar peaks in the orthologous *K*_*S*_ distributions if all the leptosporangiate species have similar synonymous substitution rates. A series of similar comparisons were performed using *Dipteris conjugata* and the above core leptosporangiates (Supplementary Figure 2).

To circumscribe the phylogenetic placements of the identified WGDs in the *K*_S_ distributions for the paranomes, we corrected the differences in synonymous substitution rates among species using two approaches. In the first approach, we first used OrthoFinder (v2.3.3) (Emms and Kelly 2019) with default settings except using “msa” for gene tree inference to identify gene families with the 16 species in Figure 1. Among the identified gene families, we selected 34 single-copy gene families to estimate the branch lengths in *K*_S_ unit using PAML (v4.9j) with the free-ratio model (Yang 2007). Then, to map all the identified *K*_S_ peaks onto the species phylogeny in the *K*_S_ unit, we halved *K*_S_ values of the identified peaks in the GMM analyses (Supplementary Figure 3) and mapped each peak from the tip toward the root of the phylogeny to date when WGD events have occurred in the phylogeny, with the assumption that duplicate genes evolved at similar substitution rates after WGD events (Figure 2d).

In the second approach, we used the software ksrates, which corrects synonymous substitution rates of other species to the rate of a focal species, i.e., the species desired to implement comparisons between the relative date of WGD and species divergence (Sensalari et al. 2021). Briefly, to identify peaks representing WGDs, ksrates fits an exponential-lognormal mixture model (ELMM) to each whole-paranome *K*_S_ distribution constructed by wgd (Zwaenepoel and Van de Peer 2018). In the ELMM model, one exponential component was used for the L-shaped SSDs (Lynch and Conery 2003) and one to five lognormal components were used for potential WGD peaks. Numbers of the lognormal components were further evaluated according to the BIC scores and the best ELMM model was plotted as shown in Supplementary Figure 5. Then, ksrates corrected synonymous substitution rates across different species. First, the divergence times among a trio of species, i.e., an outgroup and two ingroup species, including the focal one, were obtained by estimating the modes of orthologous *K*_S_ distributions with 200 bootstraps, respectively. The relative rate tests were employed to calculate *K*_S_ distances for the lineages leading to the two ingroup species after their divergence. Lastly, the divergence between the two ingroup species was rescaled by double the branch length of the focal species. After a series of calculations for each divergence event of the focal species, the rescaled divergence times were comparable with the paranome *K*_S_ distribution of the focal species. The maximum sets of trios selected to correct each divergent species pair were set as 14 and the consensus peak for multiple outgroups was set as the mean among outgroups in ksrates. Other parameters were set as default for rate correction using ksrates.

### Probabilistic Gene tree – Species Tree Reconciliation

We performed statistical gene tree – species tree reconciliation analyses using Whale (Zwaenepoel and Van de Peer 2019). We retrieved a species tree with divergence times from TimeTree (Kumar et al. 2017) (Supplementary Figure 7). We used OrthoFinder (v2.3.3) (Emms and Kelly 2019) to identify gene families with the 16 fern species using default settings except using “msa” as the method for gene tree inference. Second, after identifying 16,305 gene families, we used “orthofilter.py” in the Whale repository (https://github.com/arzwa/Whale.jl) to filter out 9,442 gene families that had no common ancestor at the root or had a large family size. Third, we used PRANK (Löytynoja 2014) to perform multiple sequence alignment of proteins in each gene family and used mrbayes (v.3.2.6) (Ronquist et al. 2012) to infer posterior probability distributions of gene trees under the LG+GAMMA model. The sample frequency was set as 10 and the number of generations was set as 110,000 to get in total 11,000 posterior samples for each gene family. Lastly, ALEobserve (Szöllõsi et al. 2013) was used to construct the conditional clade distribution containing marginal clade frequencies with a burn-in of 1,000 for each of the 6,863 gene family.

Using Whale, we carried out the probabilistic gene tree – species tree reconciliation and tested the occurrences of eight WGDs (Supplementary Figure 7) under the so-called DL+WGD model, which considers both gene duplication and loss for both SSDs and WGDs. Two DL+WGD models were adopted to incorporate various DL rates of SSDs across the species tree. In the critical branch-specific DL+WGD model, where we assumed the duplication (*λ*) and loss rates (*μ*) to be equal on each branch, a *Beta*(3,1) prior distribution was used for *η*, the parameter of the geometric prior distribution on the number of genes at the root. We used an improper flat prior for the mean branch rate *r*. The branch rates were assumed to follow a multivariate Gaussian prior with an exponential prior with mean 0.1 for the standard deviation. For the more flexible DL+WGD model with branch-specific duplication and loss rates model, we assumed an independent bivariate normal prior with mean 0 and standard deviation 1 for the mean log-duplication and loss rate, assumed a *Uniform*(−1,1) prior for the correlation coefficient of duplication and loss rate for each individual branch and assumed an exponential prior with mean 1 for the standard deviation of the branch rates.

### Collinear Analysis of Available Fern Genomes

We used i-ADHoRe (v.3.0.01) (Proost et al. 2011) to delineate both intra- and intergenomic collinear analyses with the genomes of *A. capillus-veneris, A. filiculoides* and *S. cucullata*. For the intragenomic comparisons, we identified 361, 414, and 375 anchor pairs – duplicate pairs retained in the collinear regions – in the genomes of *A. capillus-veneris, A. filiculoides*, and *S. cucullata*, respectively. After constructing *K*_S_ distributions for anchor pairs by the CODEML in the PAML package (v.4.9j) (Yang 2007), we found a larger fractions of anchor pairs with *K*_S_ values ranging from 0 to 0.1 in the genomes of *A. filiculoides* and *S. cucullata* (Supplementary Figures 8 and 9), compared to those in the genome of *A. capillus-veneris* (Supplementary Figure 10). A further look into the anchor pairs with small *K*_S_ values shows that most of them are located on short scaffolds in the assemblies of *A. filiculoides* and *S. cucullata* (Supplementary Figure 11), reflecting that the two heterosporous fern genome assemblies are still fragmental to a certain extent. These short scaffolds with fewer than ten genes and only containing anchor pairs with *K*_S_ values less than 0.1 were removed from the subsequent intergenomic analysis. In the intergenomic comparisons, an all-against-all BLASTP (v.2.6.0+) (Camacho et al. 2009) was first conducted for all the protein sequences of the three ferns with an *E*-value cut-off of 1 × 10^−5^ and ‘-max_target_seqs = 100000’. Homologous pairs were filtered based on the c-score of 0.5 as previously described (Putnam Nicholas et al. 2007) and were then fed into i-ADHoRe (v.3.0.01) (Proost et al. 2011) to estimate the collinear ratio between the three fern species (Supplementary Figure 12).

In addition, we also performed gene tree – species tree reconciliations for gene families having anchor pairs with Whale without any hypothetical WGD events, estimating the expected number of DL events on each branch by assuming an independent Exponential prior with mean 1 for the duplication and loss distances associated with each species tree branch, assuming a fixed number of 0.75 based on the average number of observed genes in a family for the *η* parameter. To reserve anchor pairs that were most likely derived from WGD events, we only kept 46, 83, and 47 anchor pairs in 179 gene families, whose *K*_S_ values fell in the ranges of [1.8,2.5], [0.6,1.2] and [1.0,1.9] according to the *K*_S_ distributions for the whole paranomes of *A. capillus-veneris, A. filiculoides*, and *S. cucullata*, respectively (Supplementary Figures 3 and 5). Then, we ended up with 137 gene families including only 30, 36, and 75 anchor pairs for *A. capillus-veneris, A. filiculoides*, and *S. cucullata*, respectively, by further removing anchor pairs that were in the gene families without a common ancestor at the root of the species tree. The posterior distributions for the probability of duplications of anchor pairs on possible branches were then depicted in Figure 4.

