## Supplementary Material for "Revisiting Ancient Polyploidy in Leptosporangiate Ferns"

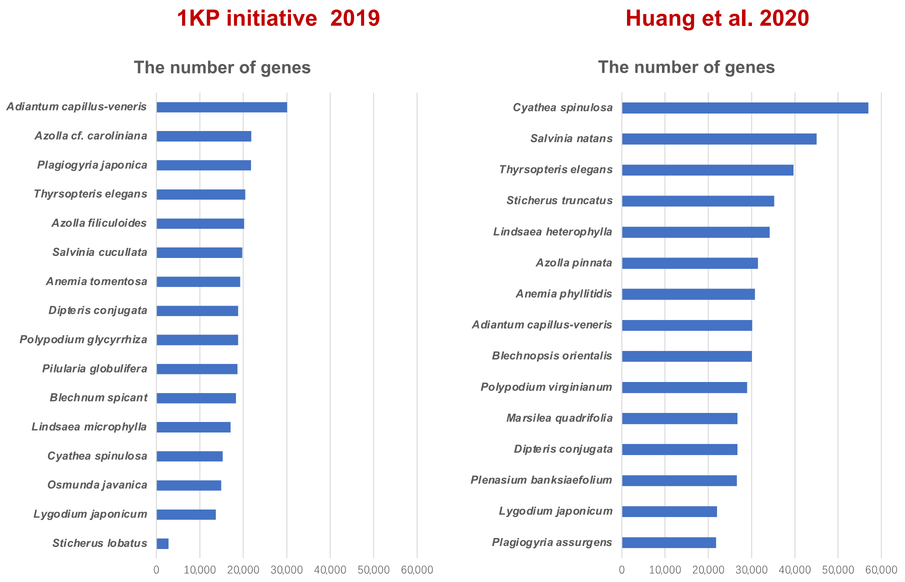


**Supplementary Figure 1.** The number of genes in the transcriptome assemblies from the 1KP initiative (2019) and Huang et al. (2020).

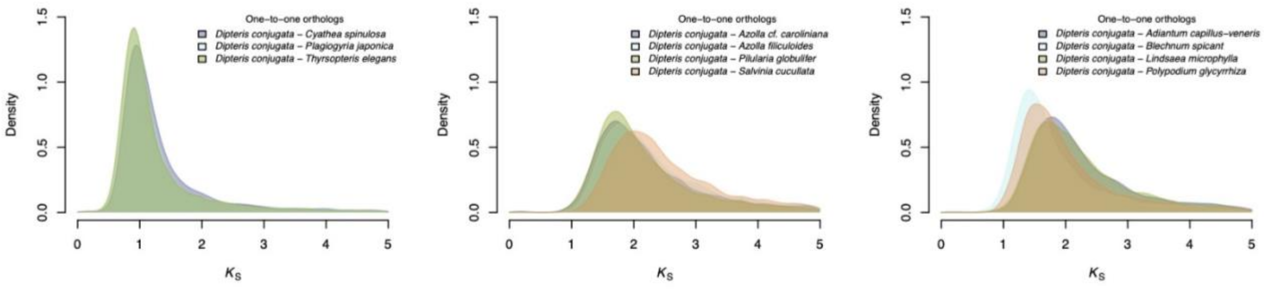


**Supplementary Figure 2.** The one-to-one orthologous $K_S$ age distributions between *Dipteris conjugata* and species from Cyatheales (left), Salviniales (middle), and Polypodiales (right).

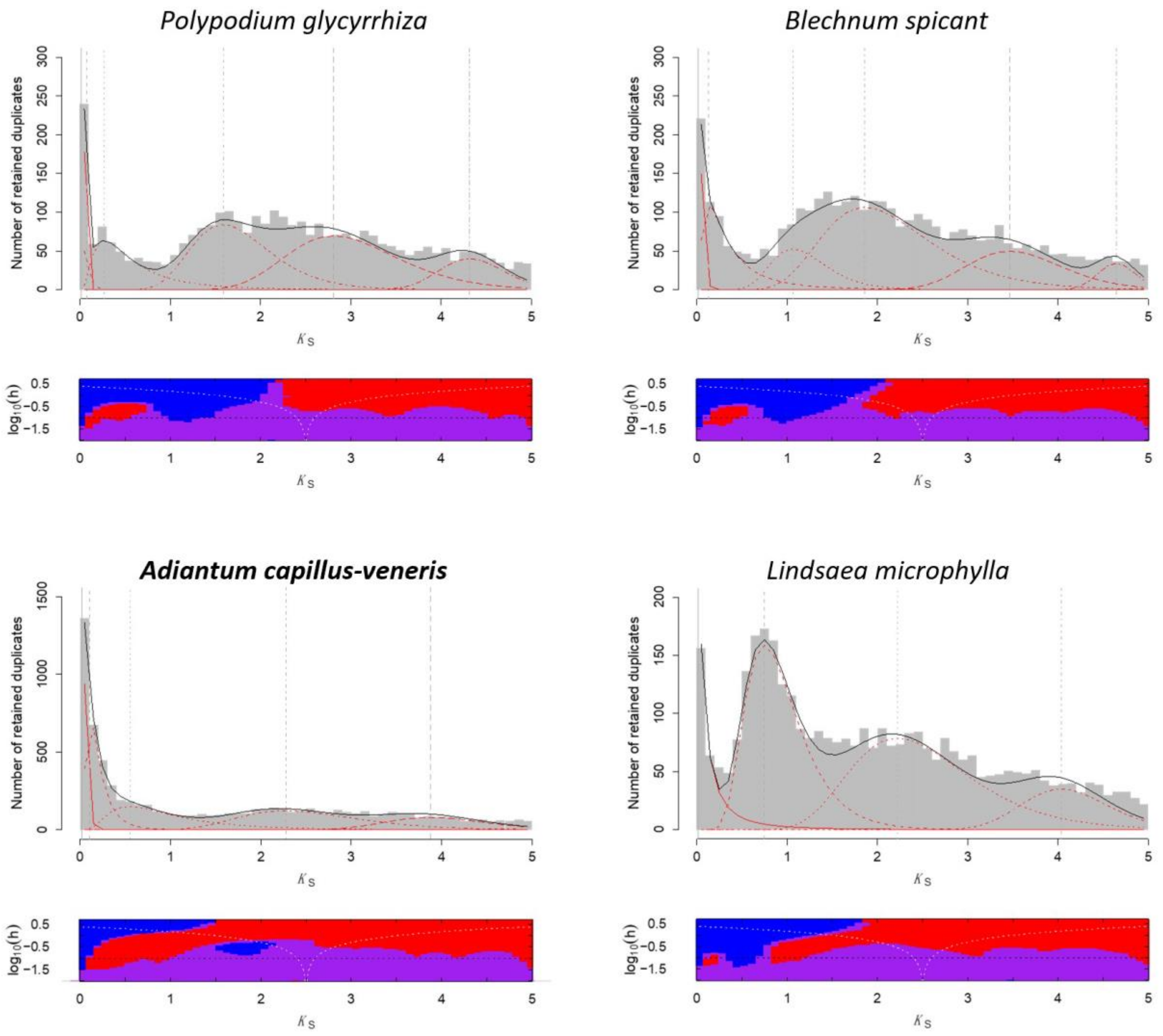


**Supplementary Figure 3.** $K_S$ distributions for the whole paranomes in different species with the Gaussian Mixture Modeling (GMM) analysis and the SiZer analysis. The optimal number of log-normal components overlaid on $K_S$ distributions in red curves with grey vertical lines representing modes, and black curves show the sum of components. In a SiZer slope plot, putative true peaks are enclosed by a blue stretch (significant upward slope) to the left and a red stretch (significant downward slope) to the right. Purple stretches correspond to no significant upward or downward slope, and gray stretches indicate regions where data is too sparse.

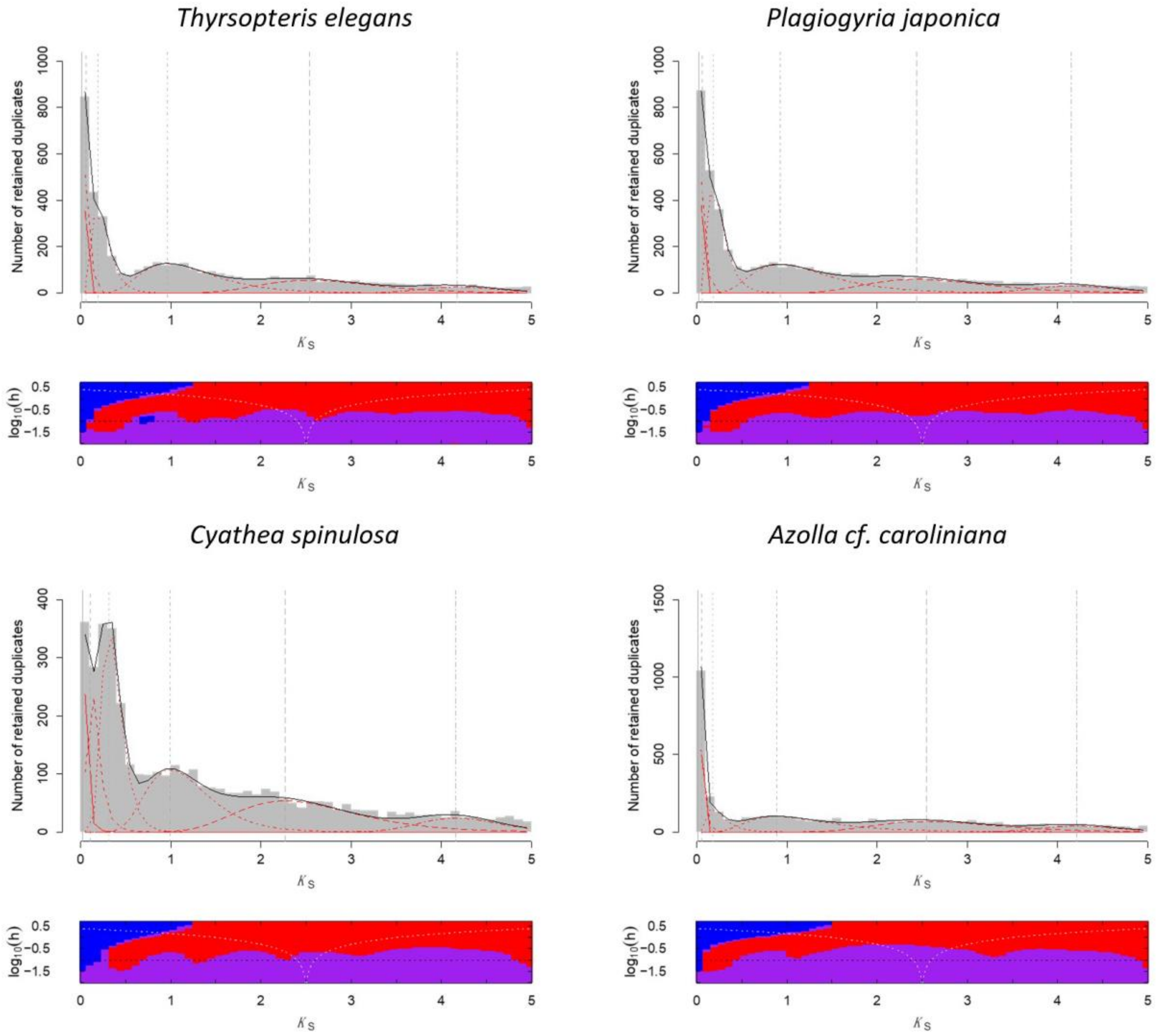


**Supplementary Figure 3 (continued).** $K_S$ distributions for the whole paranomes in different species with the Gaussian Mixture Modeling (GMM) analysis and the SiZer analysis. The optimal number of log-normal components overlaid on $K_S$ distributions in red curves with grey vertical lines representing modes, and black curves show the sum of components. In a SiZer slope plot, putative true peaks are enclosed by a blue stretch (significant upward slope) to the left and a red stretch (significant downward slope) to the right. Purple stretches correspond to no significant upward or downward slope, and gray stretches indicate regions where data is too sparse.

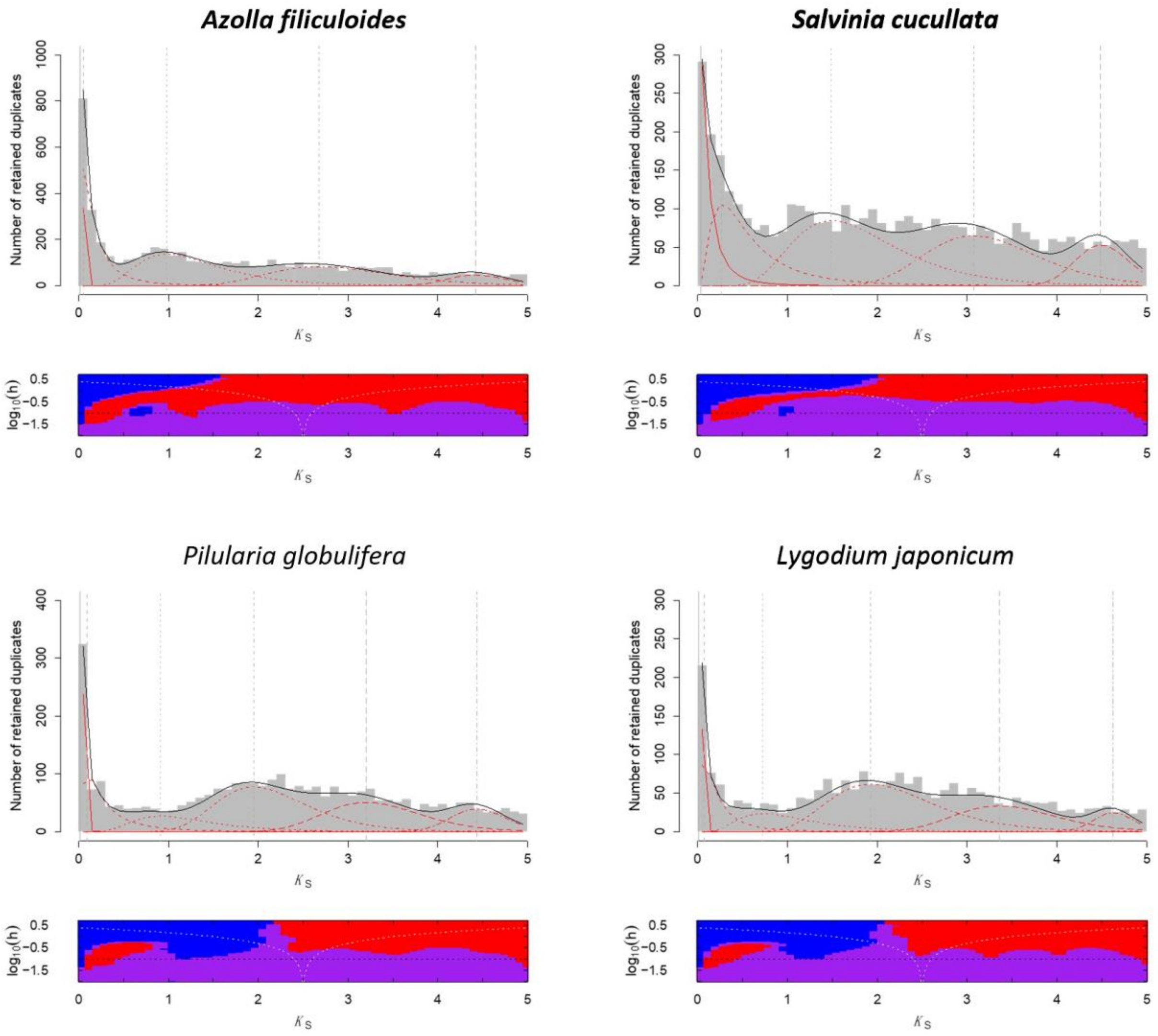


**Supplementary Figure 3 (continued).** $K_S$ distributions for the whole paranomes in different species with the Gaussian Mixture Modeling (GMM) analysis and the SiZer analysis. The optimal number of log-normal components overlaid on $K_S$ distributions in red curves with grey vertical lines representing modes, and black curves show the sum of components. In a SiZer slope plot, putative true peaks are enclosed by a blue stretch (significant upward slope) to the left and a red stretch (significant downward slope) to the right. Purple stretches correspond to no significant upward or downward slope, and gray stretches indicate regions where data is too sparse.

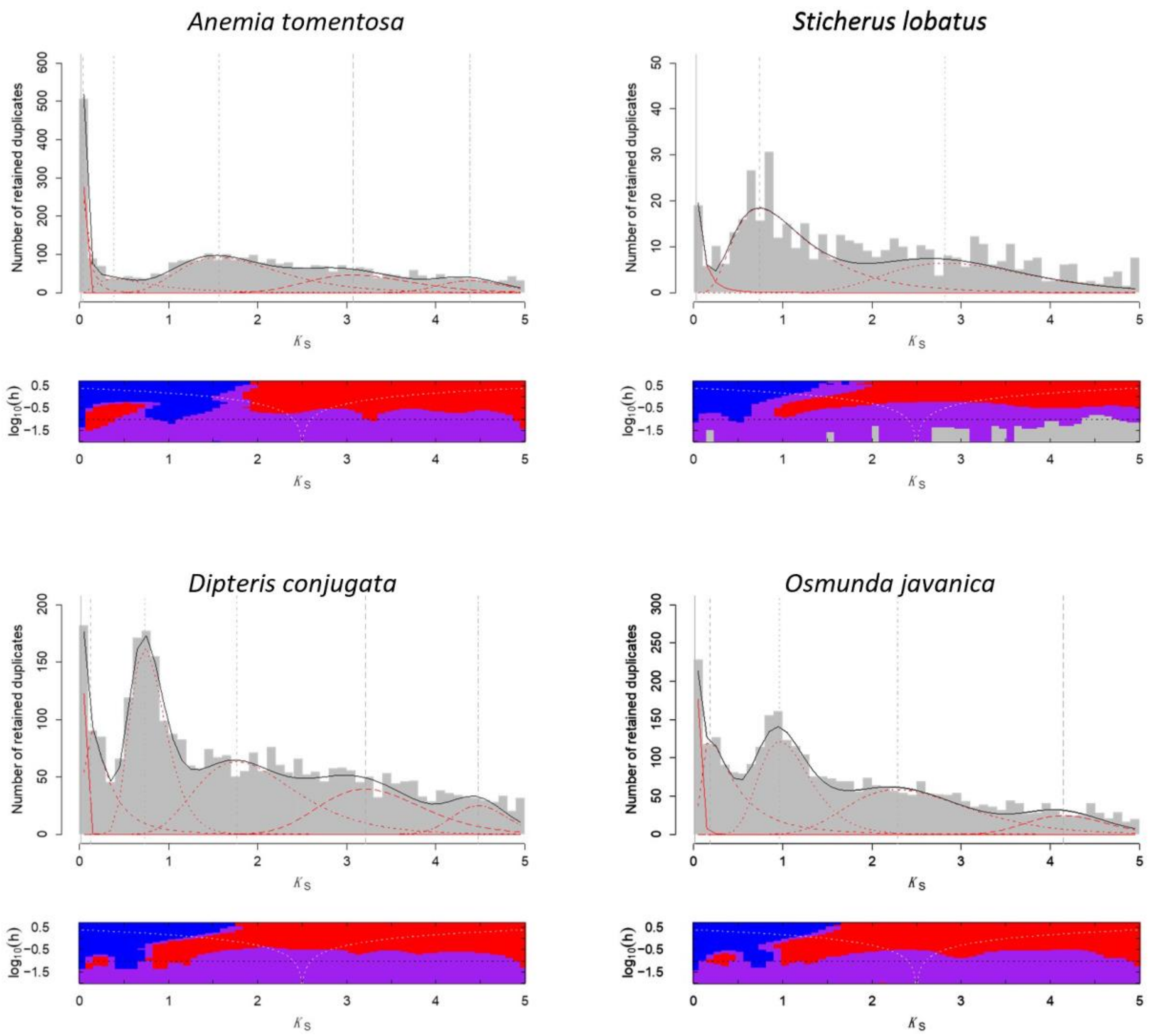


**Supplementary Figure 3 (continued).** $K_S$ distributions for the whole paranomes in different species with the Gaussian Mixture Modeling (GMM) analysis and the SiZer analysis. The optimal number of log-normal components overlaid on $K_S$ distributions in red curves with grey vertical lines representing modes, and black curves show the sum of components. In a SiZer slope plot, putative true peaks are enclosed by a blue stretch (significant upward slope) to the left and a red stretch (significant downward slope) to the right. Purple stretches correspond to no significant upward or downward slope, and gray stretches indicate regions where data is too sparse.

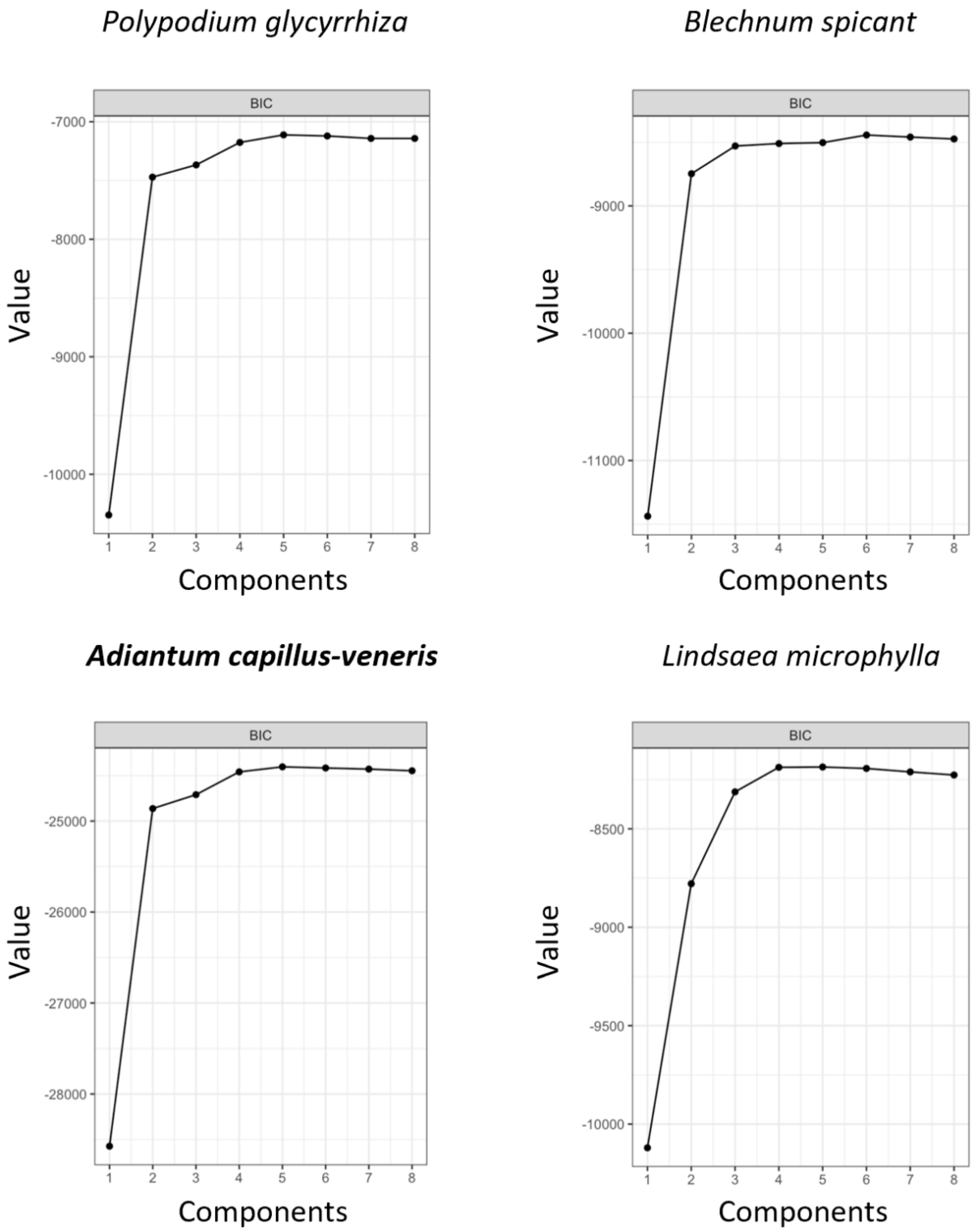


**Supplementary Figure 4.** The Bayesian Information Criterion (BIC) score in the Gaussian mixture modeling analysis for different species in Supplementary Figure 3.

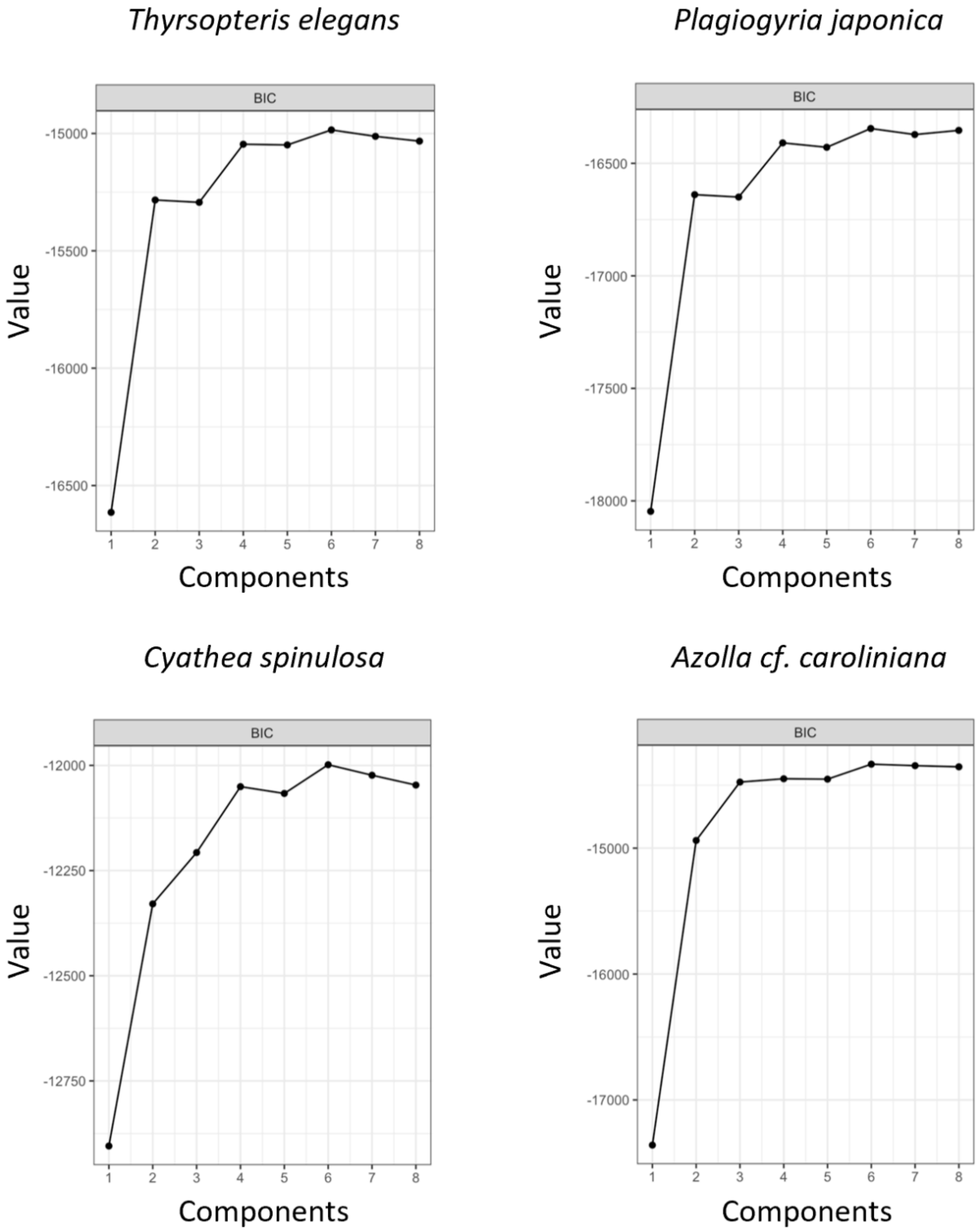


**Supplementary Figure 4 (continued).** The Bayesian Information Criterion (BIC) score in the Gaussian mixture modeling analysis for different species in Supplementary Figure 3.

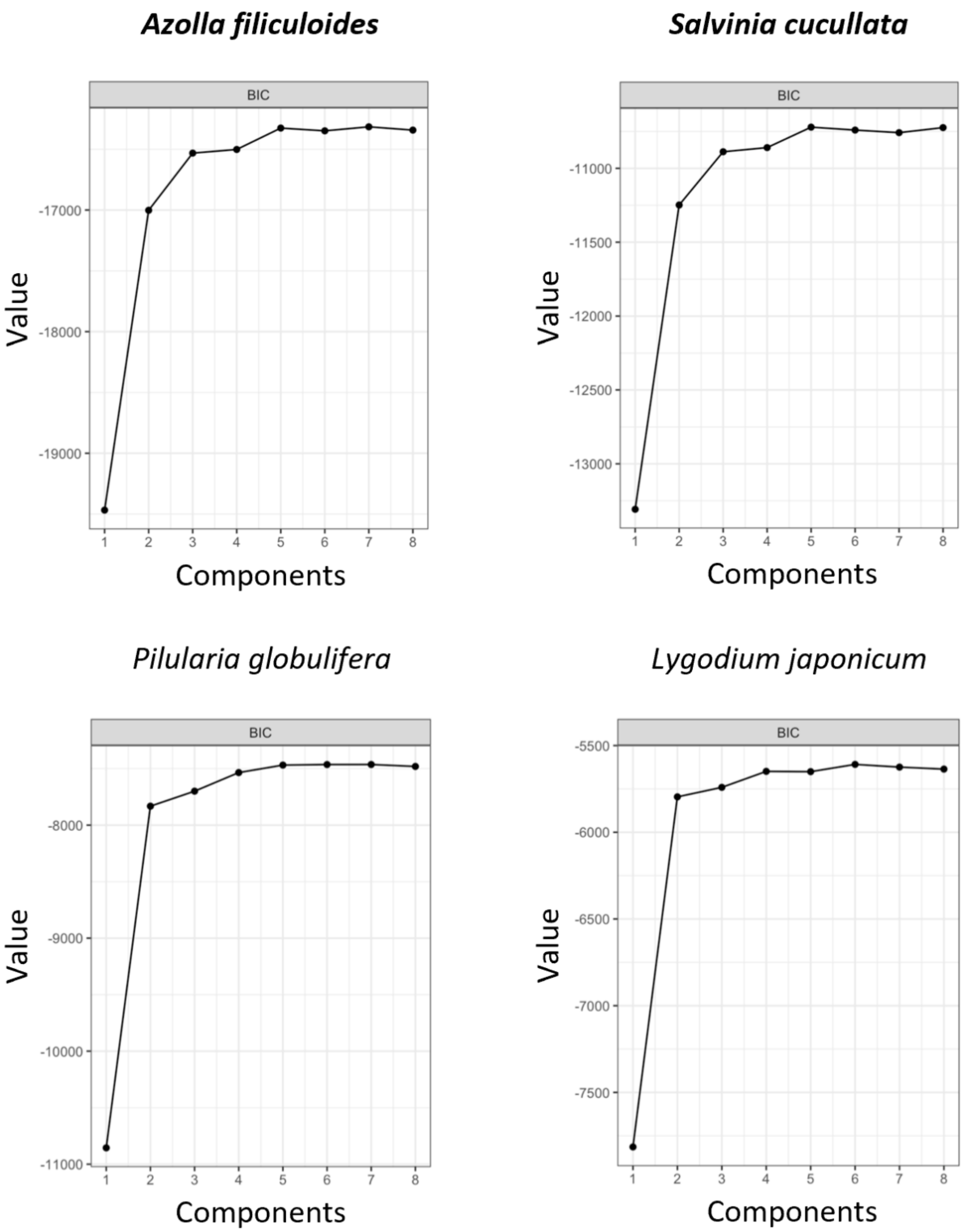


**Supplementary Figure 4 (continued).** The Bayesian Information Criterion (BIC) score in the Gaussian mixture modeling analysis for different species in Supplementary Figure 3.

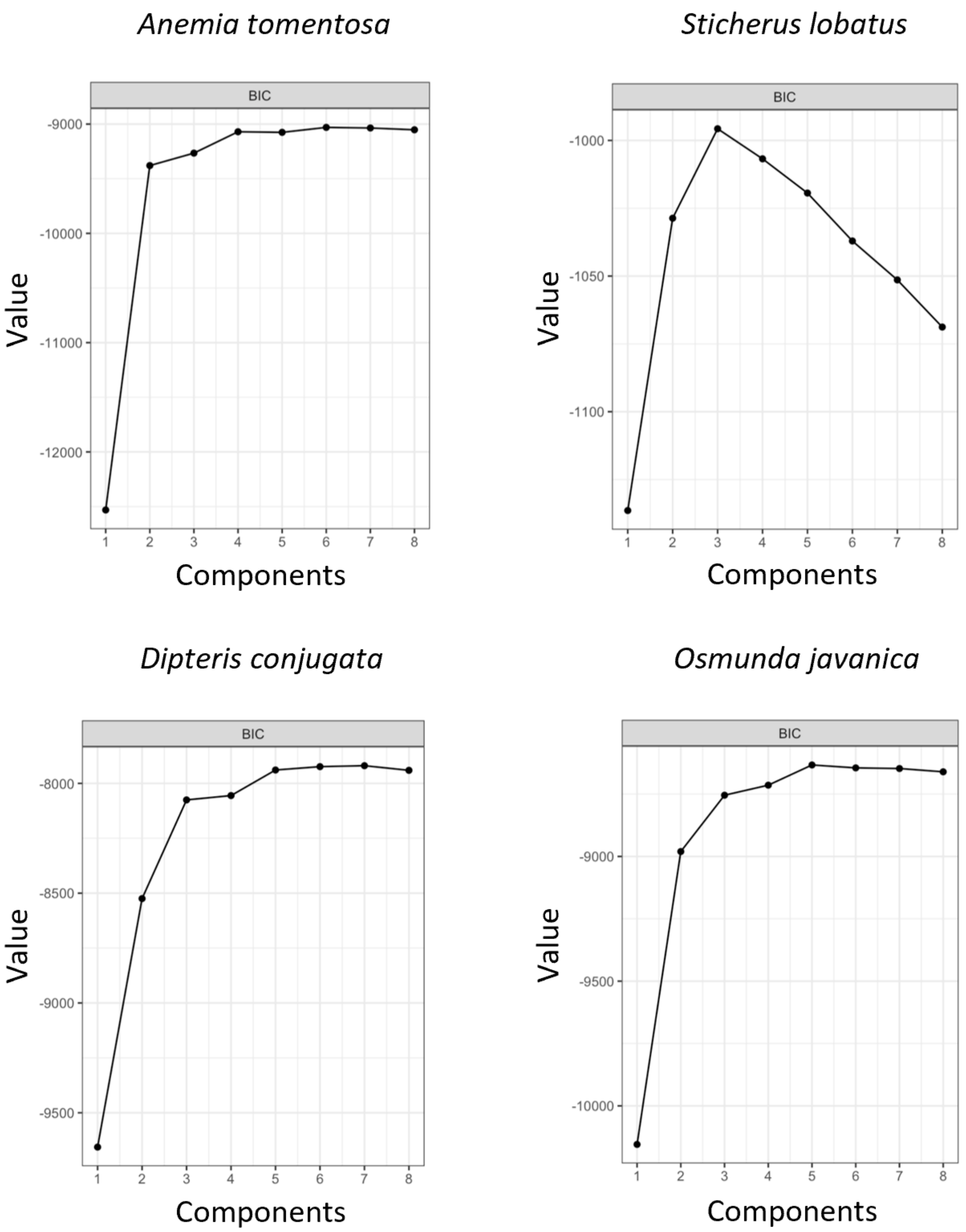


**Supplementary Figure 4 (continued).** The Bayesian Information Criterion (BIC) score in the Gaussian mixture modeling analysis for different species in Supplementary Figure 3.

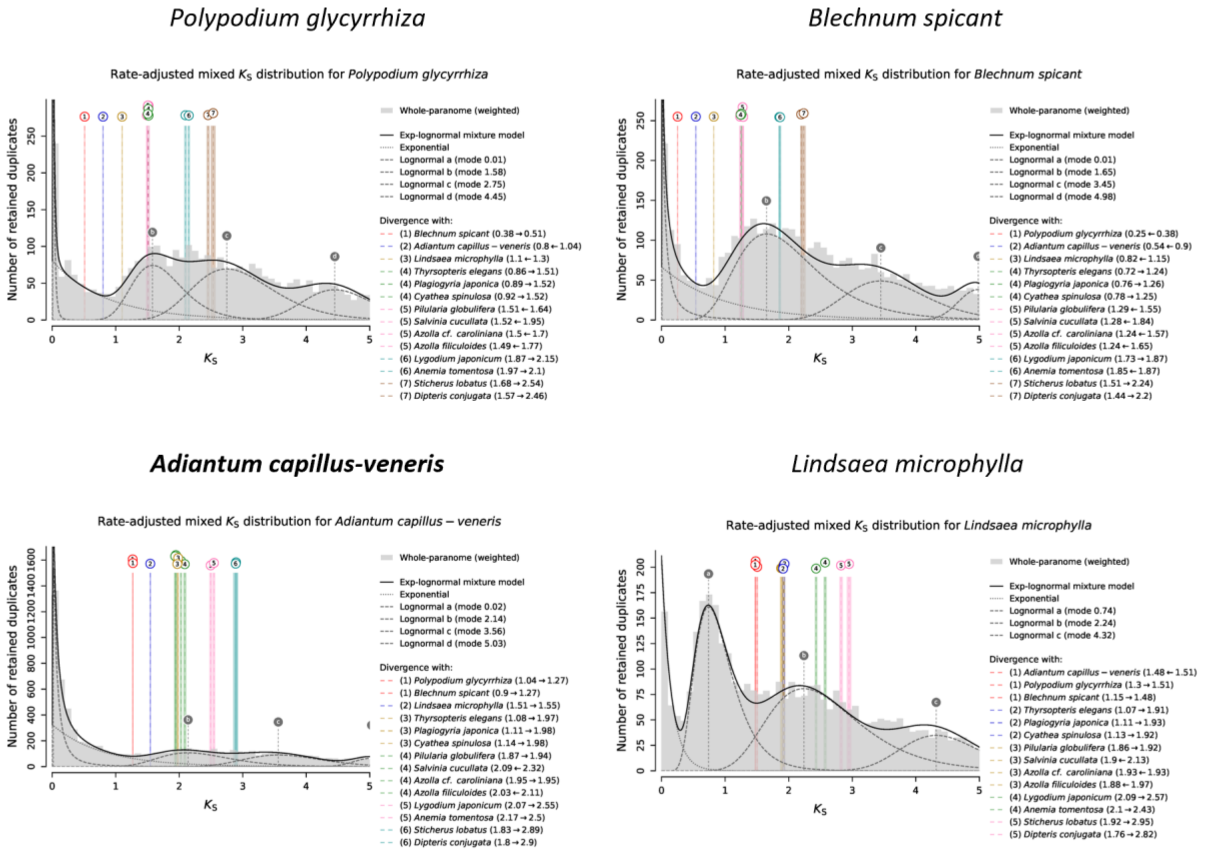


**Supplementary Figure 5.** The analyses of ksrates for different species. $K_S$ distributions for the whole paranomes of different species are overlaid with rate-adjusted species events in colored vertical lines. The overall mixture model in the dark solid line of each paralogous $K_S$ distribution consists of an exponential component in dotted gray curve and optimized log-normal components in dashed gray curves. Each log-normal component is labeled with a letter, shown as vertical dashed gray lines with circular labels. Rate-adjusted mode estimates of orthologous $K_S$ distributions between a focal species and other species, representing speciation events, are drawn as numbered vertical long-dashed lines, with associated colored boxes showing the standard deviation and the mean of estimated mode. Lines representing the same speciation event in the phylogeny share color and numbering. Horizontal arrows in figure legends indicate the $K_S$ shifts produced by the substitution rate adjustments.

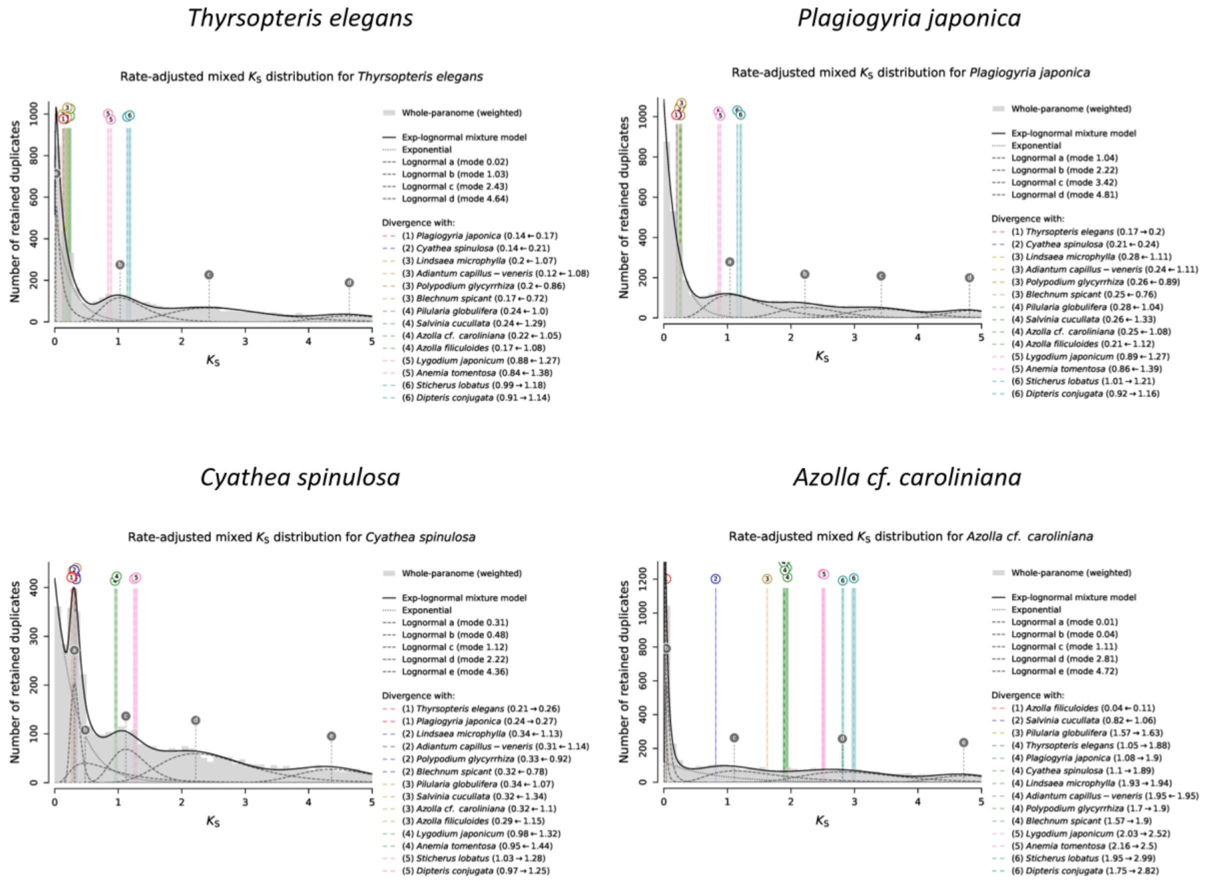


**Supplementary Figure 5 (continued).** The analyses of ksrates for different species. $K_S$ distributions for the whole paranomes of different species are overlaid with rate-adjusted species events in colored vertical lines. The overall mixture model in the dark solid line of each paralogous $K_S$ distribution consists of an exponential component in dotted gray curve and optimized log-normal components in dashed gray curves. Each log-normal component is labeled with a letter, shown as vertical dashed gray lines with circular labels. Rate-adjusted mode estimates of orthologous $K_S$ distributions between a focal species and other species, representing speciation events, are drawn as numbered vertical long-dashed lines, with associated colored boxes showing the standard deviation and the mean of estimated mode. Lines representing the same speciation event in the phylogeny share color and numbering. Horizontal arrows in figure legends indicate the $K_S$ shifts produced by the substitution rate adjustments.

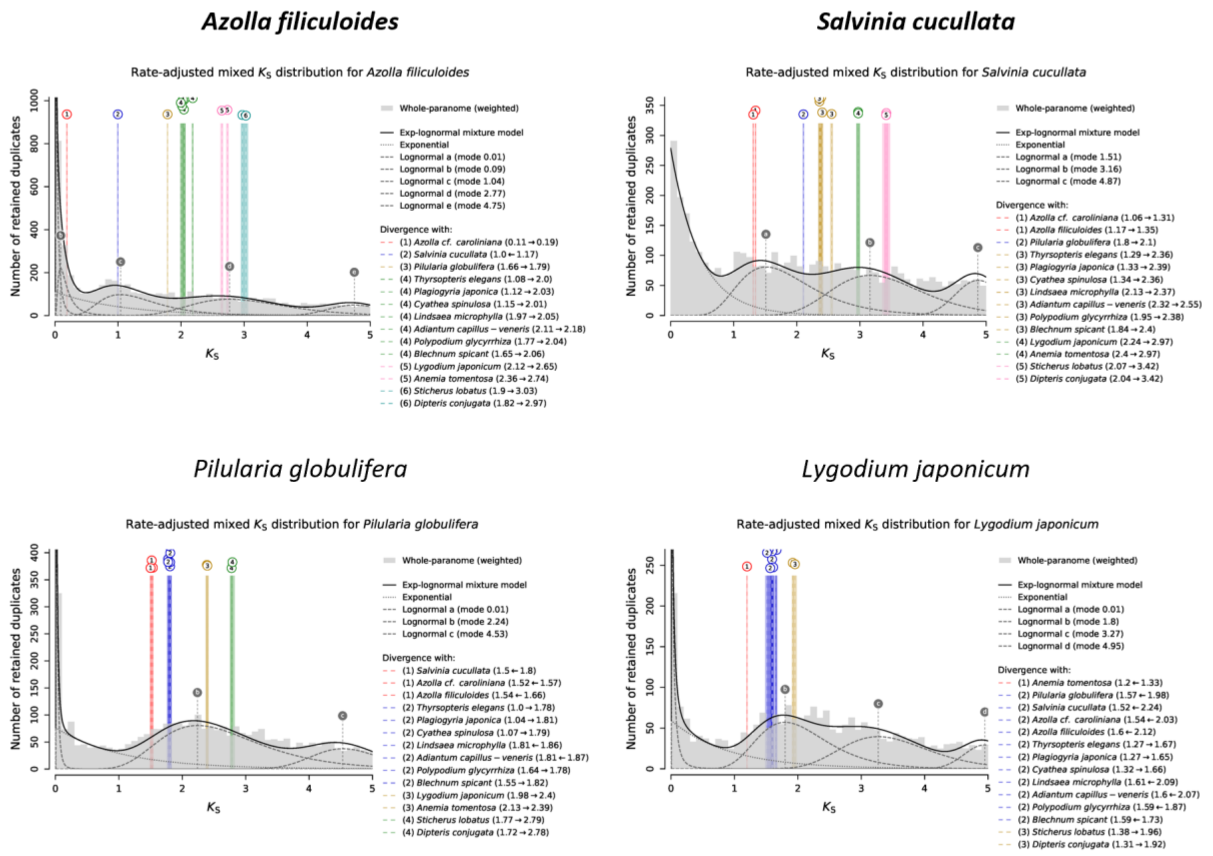


**Supplementary Figure 5 (continued).** The analyses of ksrates for different species. $K_S$ distributions for the whole paranomes of different species are overlaid with rate-adjusted species events in colored vertical lines. The overall mixture model in the dark solid line of each paralogous $K_S$ distribution consists of an exponential component in dotted gray curve and optimized log-normal components in dashed gray curves. Each log-normal component is labeled with a letter, shown as vertical dashed gray lines with circular labels. Rate-adjusted mode estimates of orthologous $K_S$ distributions between a focal species and other species, representing speciation events, are drawn as numbered vertical long-dashed lines, with associated colored boxes showing the standard deviation and the mean of estimated mode. Lines representing the same speciation event in the phylogeny share color and numbering. Horizontal arrows in figure legends indicate the $K_S$ shifts produced by the substitution rate adjustments.

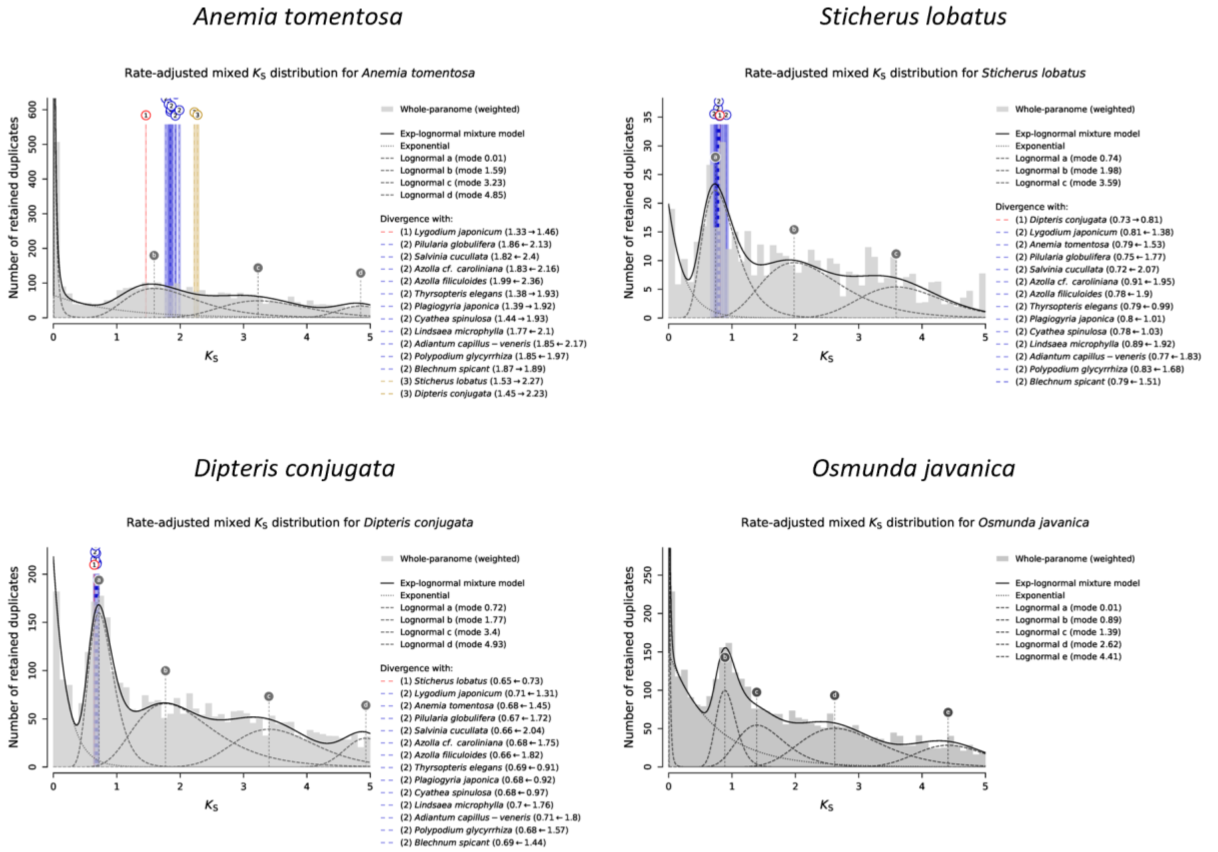


**Supplementary Figure 5 (continued).** The analyses of ksrates for different species. $K_S$ distributions for the whole paranomes of different species are overlaid with rate-adjusted species events in colored vertical lines. The overall mixture model in the dark solid line of each paralogous $K_S$ distribution consists of an exponential component in dotted gray curve and optimized log-normal components in dashed gray curves. Each log-normal component is labeled with a letter, shown as vertical dashed gray lines with circular labels. Rate-adjusted mode estimates of orthologous $K_S$ distributions between a focal species and other species, representing speciation events, are drawn as numbered vertical long-dashed lines, with associated colored boxes showing the standard deviation and the mean of estimated mode. Lines representing the same speciation event in the phylogeny share color and numbering. Horizontal arrows in figure legends indicate the $K_S$ shifts produced by the substitution rate adjustments.

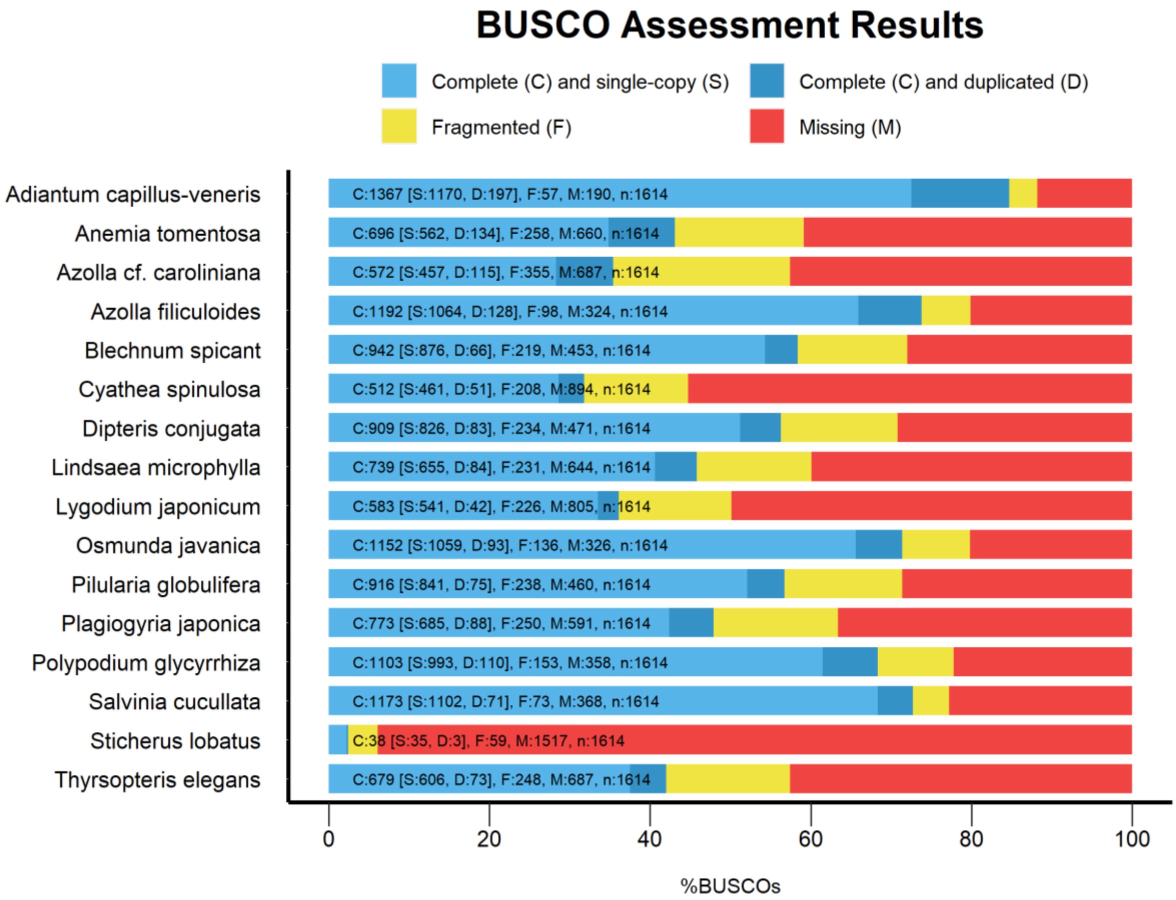


**Supplementary Figure 6.** BUSCO analysis for the transcriptome assemblies from the 1KP initiative (2019).

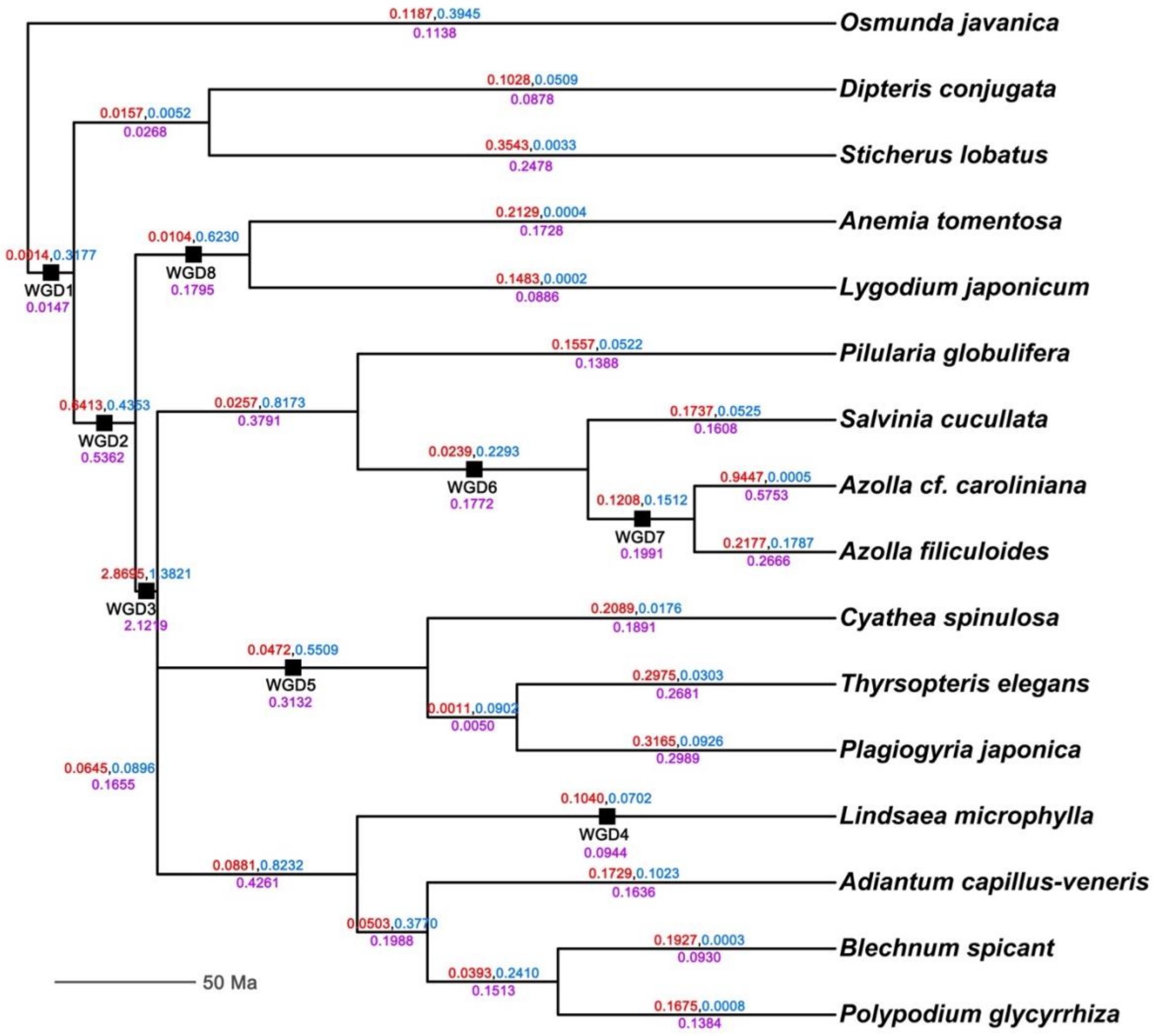


**Supplementary Figure 7.** The time-calibrated species trees from TimeTree. The numbers above a branch are the duplication (red) and loss (blue) rates in the relaxed branch-specific model. The number below a branch is the duplication and loss rates (purple), which are equal in the critical branch-specific model. The black squares on branches are the eight WGDs that were tested in the DL+WGD model.

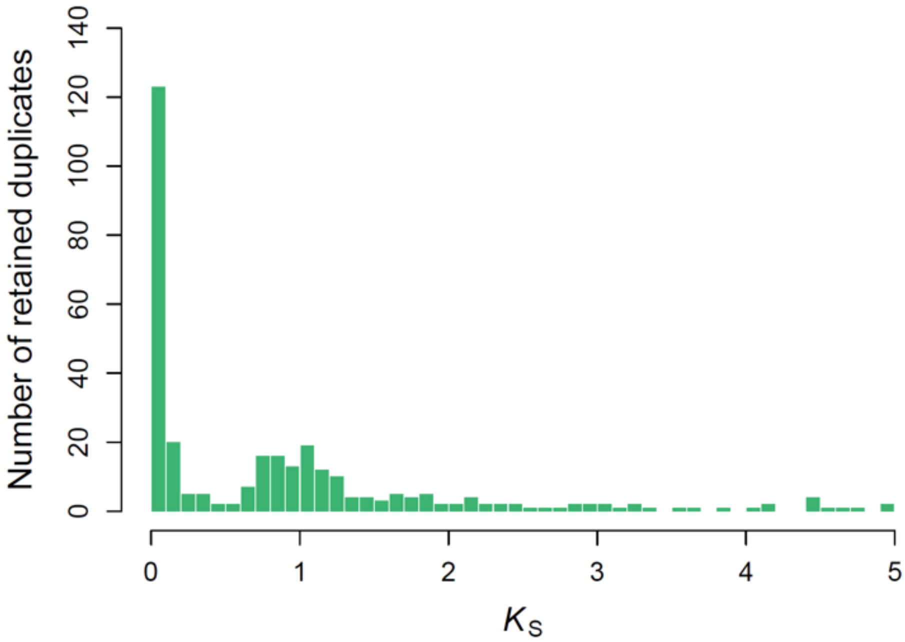


**Supplementary Figure 8.** $K_S$ distributions for the anchor pairs identified in *Azolla filiculoides*. Anchor pairs with $K_S$ values less than 0.1 tend to be located on short scaffolds in the genome assembly (see Methods and Supplementary Figure 11)

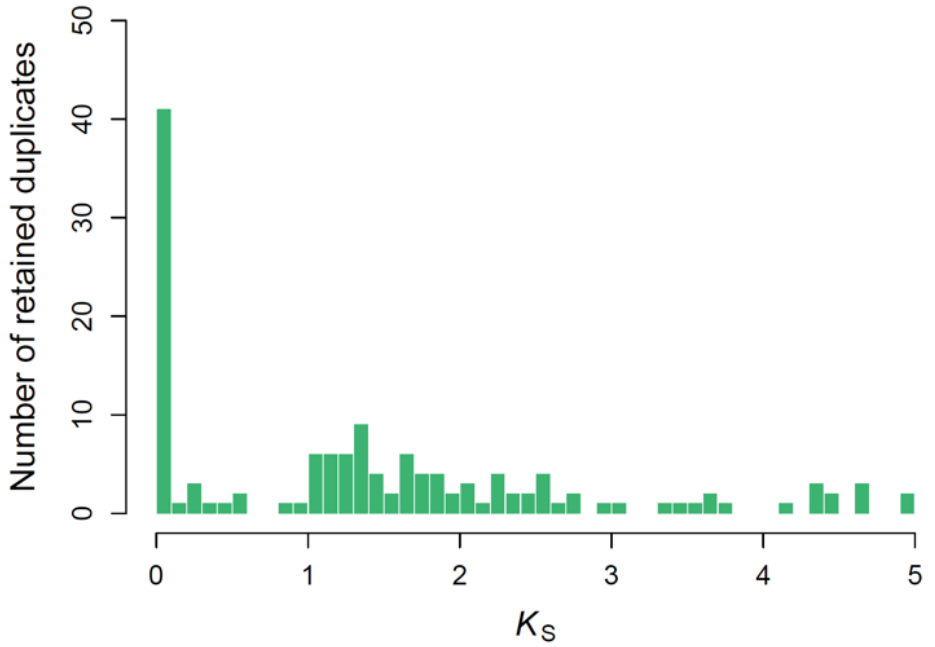


**Supplementary Figure 9.** $K_S$ distributions for the anchor pairs identified in *Salvinia cucullata*. Anchor pairs with $K_S$ values less than 0.1 tend to be located on short scaffolds in the genome assembly (see Methods and Supplementary Figure 11).

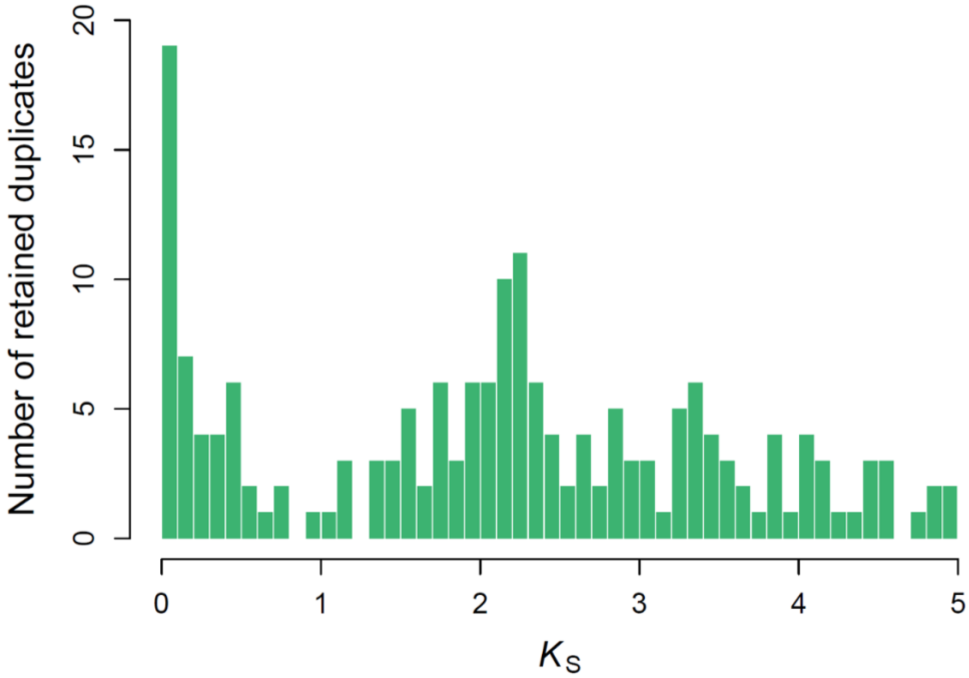


**Supplementary Figure 10.** $K_S$ distributions for the anchor pairs identified in *Adiantum capillus-veneris*.

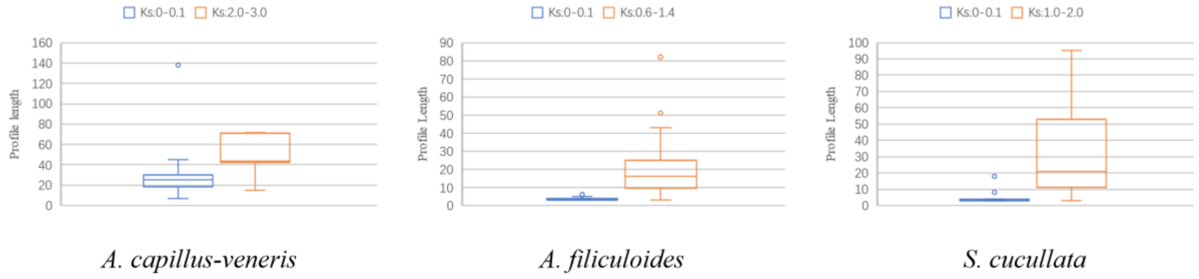


**Supplementary Figure 11.** Box plots of the number of genes without tandem duplicates (profile lengths) on scaffolds having anchor pairs with $K_S$ values less than 0.1 and those having anchor pairs with $K_S$ values near a potential WGD peak in the three fern genomes.

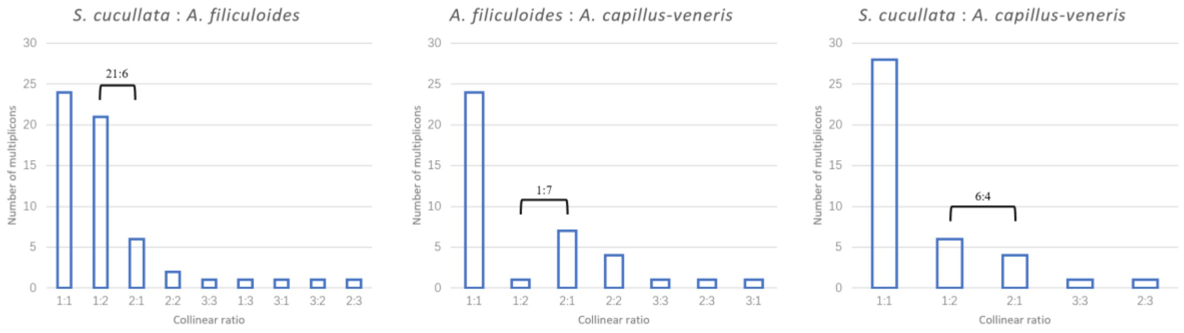


**Supplementary Figure 12.** Ratios of collinear blocks for pairwise intergenomic comparisons among the three genome-available ferns (see Methods). Collinear blocks with at least two species were retrieved and counted for the ratios, which turn out to be 2 : 1 : 1 for *A. filiculoides* : *S. cucullata* : *A. capillus-veneris*. The number above bars show the number of collinear blocks for a certain collinear ratio.

**Supplementary Table 1. Taxonomy, number of genes/unigenes and data source of fern species involved in this study.**

| Clade | Order | Family | Species | Number of genes | Source of data |
|---|---|---|---|---|---|
| Core Leptosporangiates | Cyatheales | Cyatheaceae | *Cyathea spinulosa* | 15288 | 1KP initiative (2019) |
| Core Leptosporangiates | Cyatheales | Thyrsopteridaceae | *Thyrsopteris elegans* | 20444 | 1KP initiative (2019) |
| Leptosporangiates | Gleicheniales | Gleicheniaceae | *Sticherus lobatus* | 2789 | 1KP initiative (2019) |
| Leptosporangiates | Gleicheniales | Dipteridaceae | *Dipteris conjugata* | 18791 | 1KP initiative (2019) |
| Leptosporangiates | Osmundales | Osmundaceae | *Osmunda javanica* | 14952 | 1KP initiative (2019) |
| Core Leptosporangiates | Cyatheales | Plagiogyriaceae | *Plagiogyria japonica* | 21769 | 1KP initiative (2019) |
| Core Leptosporangiates | Polypodiales | Polypodiaceae | *Polypodium glycyrrhiza* | 18785 | 1KP initiative (2019) |
| Core Leptosporangiates | Polypodiales | Blechnaceae | *Blechnum spicant* | 18308 | 1KP initiative (2019) |
| Core Leptosporangiates | Polypodiales | Lindsaeaceae | *Lindsaea microphylla* | 17074 | 1KP initiative (2019) |
| Core Leptosporangiates | Salviniales | Salviniaceae | *Azolla cf. caroliniana* | 21852 | 1KP initiative (2019) |
| Core Leptosporangiates | Salviniales | Marsileaceae | *Pilularia globulifera* | 18655 | 1KP initiative (2019) |
| Leptosporangiates | Schizaeales | Anemiaceae | *Anemia tomentosa* | 19274 | 1KP initiative (2019) |
| Leptosporangiates | Schizaeales | Lygodiaceae | *Lygodium japonicum* | 13716 | 1KP initiative (2019) |
| Core Leptosporangiates | Cyatheales | Cyatheaceae | *Cyathea spinulosa* | 56985 | Huang et al. (2020) |
| Core Leptosporangiates | Cyatheales | Thyrsopteridaceae | *Thyrsopteris elegans* | 39658 | Huang et al. (2020) |
| Leptosporangiates | Gleicheniales | Gleicheniaceae | *Sticherus truncatus* | 35205 | Huang et al. (2020) |
| Leptosporangiates | Gleicheniales | Dipteridaceae | *Dipteris conjugata* | 26737 | Huang et al. (2020) |
| Leptosporangiates | Osmundales | Osmundaceae | *Plenasium banksiaefolium* | 26583 | Huang et al. (2020) |
| Core Leptosporangiates | Cyatheales | Plagiogyriaceae | *Plagiogyria assurgens* | 21829 | Huang et al. (2020) |
| Core Leptosporangiates | Polypodiales | Polypodiaceae | *Polypodium virginianum* | 28943 | Huang et al. (2020) |
| Core Leptosporangiates | Polypodiales | Blechnaceae | *Blechnopsis orientalis* | 30101 | Huang et al. (2020) |
| Core Leptosporangiates | Polypodiales | Lindsaeaceae | *Lindsaea heterophylla* | 34154 | Huang et al. (2020) |

| | | | | | |
|---|---|---|---|---|---|
| Core Leptosporangiates | Salviniales | Salviniaceae | *Salvinia natans* | 45028 | Huang et al. (2020) |
| Core Leptosporangiates | Salviniales | Salviniaceae | *Azolla pinnata* | 31501 | Huang et al. (2020) |
| Core Leptosporangiates | Salviniales | Marsileaceae | *Marsilea quadrifolia* | 26751 | Huang et al. (2020) |
| Leptosporangiates | Schizaeales | Anemiaceae | *Anemia phyllitidis* | 30786 | Huang et al. (2020) |
| Leptosporangiates | Schizaeales | Lygodiaceae | *Lygodium japonicum* | 21975 | Huang et al. (2020) |
| Core Leptosporangiates | Salviniales | Salviniaceae | *Salvinia cucullata* | 19780 | Li et al. (2018) |
| Core Leptosporangiates | Salviniales | Salviniaceae | *Azolla filiculoides* | 20203 | Li et al. (2018) |
| Core Leptosporangiates | Polypodiales | Pteridaceae | *Adiantum capillus-veneris* | 31244 | Under review |
